# Placental *Igf1* Overexpression Sex-Specifically Impacts Mouse Placenta Structure, Altering Offspring Striatal Development and Behavior

**DOI:** 10.1101/2025.03.27.644829

**Authors:** Annemarie J. Carver, Faith M. Fairbairn, Robert J. Taylor, Shanmukh Boggarapu, Njenga R. Kamau, Amrita Gajmer, Hanna E. Stevens

**Affiliations:** Interdisciplinary Graduate Program in Genetics, University of Iowa, IA, USA; Department of Psychiatry, Carver College of Medicine, University of Iowa, IA, USA; Iowa Neuroscience Institute, Carver College of Medicine, University of Iowa, IA, USA; Hawk-Intellectual and Developmental Disabilities Research Center, University of Iowa, IA, USA

**Keywords:** Placenta, Neurodevelopment, Neurodevelopmental Disorders, Insulin-like Growth Factor 1, Neuroplacentology, Striatum, Ganglionic Eminence, CRISPR

## Abstract

Insulin-like growth factor 1 (IGF1) is produced primarily in the placenta *in utero* and is an essential hormone for neurodevelopment. Specifically, how placental IGF1 production persistently influences the brain is unclear. This study evaluated the effects of placental *Igf1* overexpression on embryonic and postnatal brain development, particularly for striatum, a region highly linked to neurodevelopmental disorders. Placental *Igf1* was overexpressed via placental-targeted CRISPR manipulation. This overexpression altered placenta structure and function distinctly in females and males. Early differences in placental function altered the trajectory of striatal development, as adult females showed persistent changes in striatal cell composition and striatal dependent behavior while males were less affected in brain and behavior outcomes. Overall, these results demonstrate that placental *Igf1* expression alters striatal development and behavior in ways relevant to neurodevelopmental disorders. These findings expand our understanding of placental influence on neurodevelopment and will aid in identifying placental-targeted preventive interventions.

**GRAPHICAL ABSTRACT:** 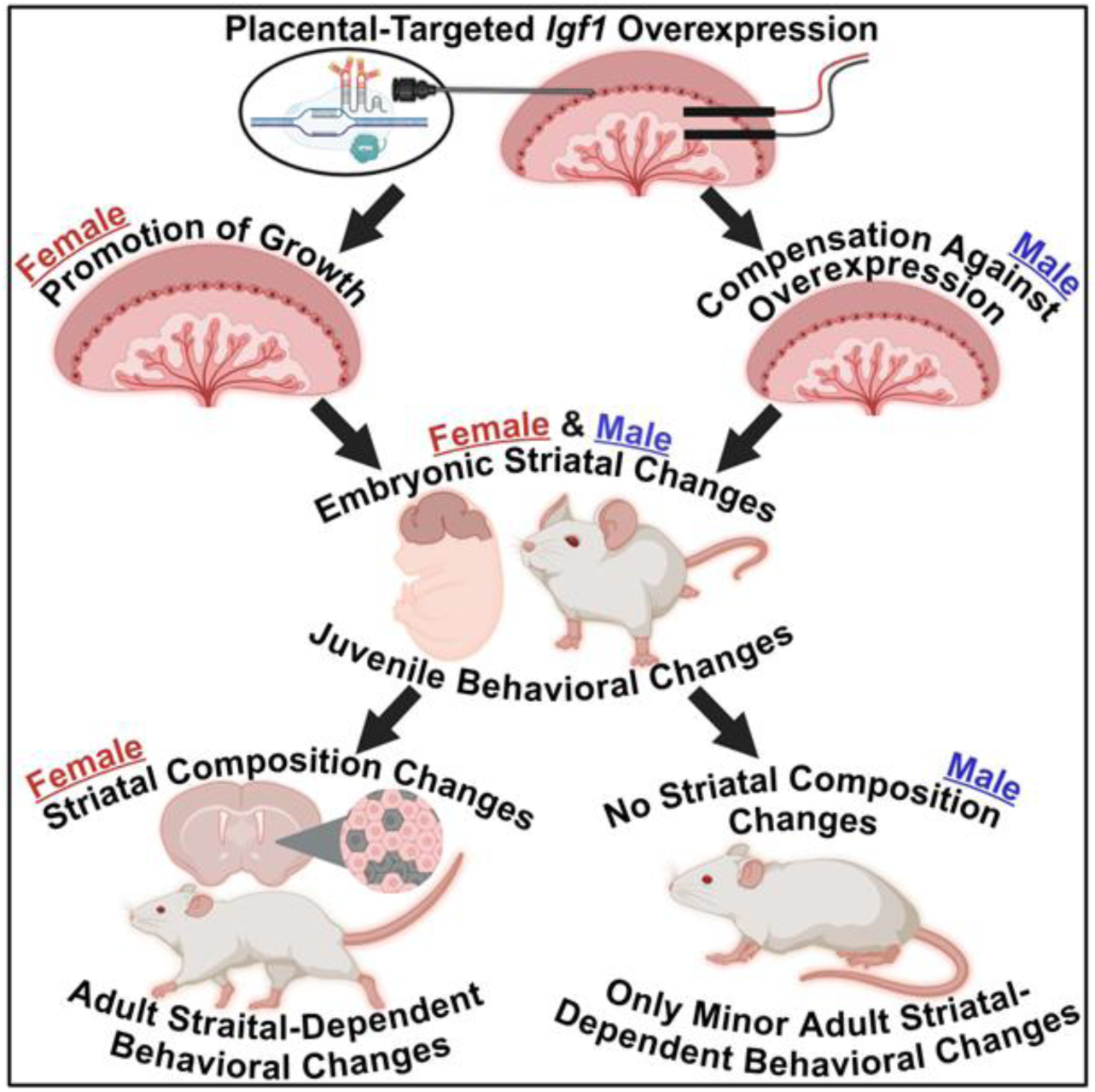

## INTRODUCTION

The placenta provides vital growth factors essential for embryonic neurodevelopment.^1,2^ Despite the known link between the placenta and neurodevelopment, the specific impact of placental factors on long-term brain structure and function has been minimally studied.^3^ In both mice and humans, the placenta is the primary source of Insulin-like Growth Factor 1 (IGF1) to the embryo and fetus prior to birth.^4,5^ IGF1 is necessary for meditating *in utero* growth generally and particularly for neurodevelopment.^4–6^ IGF1 is known to promote the proliferation, differentiation, and maturation of many cells in the brain during prenatal development, including neurons, astrocytes, and oligodendrocytes.^4,6,7^ Specifically, IGF1 promotes neuronal precursors to preferentially differentiate into neurons.^8^ Additionally, overexpression of *Igf1* within brain cells causes an enlargement of specific brain regions as well as total brain weight^6,9,10^ In particular, exogenous IGF1 stimulates cell cycle transition by promoting the transition from G1 to S phase of neural stem cells within the cerebral cortex.^11^ IGF1 activates these processes through its actions via its receptor, IGF1R.^10,12^ The receptor IGF1R is widely expressed throughout the developing brain and its signaling pathway is crucial for proper brain growth.^6^ Current research highlights the strong influence of IGF1 signaling on neurodevelopment, but links with its placental source are less understood.

Previous studies in mice have shown that the development of the striatum may be susceptible to changes in IGF signaling. This includes increased volume and neuron number in the caudate putamen (a subregion of the striatum)^9^ after overexpression of *Igf1* in the developing brain and reduced volume and parvalbumin-containing neurons of the striatum after a complete knockout of *Igf1*.^6,13^ During *in utero* neurodevelopment, the embryonic ganglionic eminence gives rise to the striatum.^14^ The striatum is a functional region within the forebrain that contributes to motor regulation and procedural and habit learning.^15–17^ These behaviors are highly associated with prevalent neurodevelopmental disorders (NDDs) including autism spectrum disorder (ASD) and attention-deficit/hyperactivity disorder (ADHD).^15,18^ Additionally, increased striatal size^15,19–23^ as well as increased IGF1 serum levels^24–28^ have been linked to both ASD and ADHD. Placental-specific manipulations have been examined for their impacts on cerebral cortical and cerebellar development,^29,30^ but little information exists about placental influence on striatal development. Prenatal brain developmental processes and the influence of factors like IGF1 are critically important to better understand, diagnose, and treat brain disorders, particularly NDDs that have an early onset. Changes to the gestational environment, including placental genetics, are likely to influence risk for NDDs; this is in line with the developmental origins of health and disease hypothesis.^31^

NDDs as well as placental anomalies more prevalently and severely impact males than females.^32–34^ A 3:1 and 2:1 male prevalence is seen in ASD and ADHD diagnosis, respectively.^35–37^ The male prevalence of NDD-associated impairment has led to a focus on predominantly male outcomes in the past. However, it is critical to understand sex differences in these disorders and in the biological mechanisms which contribute to NDDs by studying both males and females. IGF1 interacts with sex hormones and has been shown to have differing effects on the brain in males and females.^38–41^ Given these various sex differences, placental IGF1 may alter neurodevelopment differently in males and females.

Outside of neurodevelopment, IGF signaling is also critical within the placenta for its own proper structure and functions which support prenatal offspring development.^5^ IGF1 signaling is regulated by IGF binding proteins that control bioavailability within the placenta and body.^42^ IGF1 promotes growth-related pathways in the placenta, including angiogenesis.^5^ IGF1 is primarily expressed in endothelial cells of the fetal region of human placenta and labyrinth zone of mouse placenta.^43,44^ In the mouse, this region is bordered by the fetal junctional zone which is adjacent to the maternal decidua; previous studies show that placental subregions influence each other’s growth and function.^45^ Additionally, the labyrinth zone is critical for nutrient and hormone transport to the offspring^1,46^ and is highly vascularized. Therefore, the regulation of angiogenesis and IGF signaling in this organ is critical for offspring development.^1,45,46^

Using an established placental-targeted CRISPR manipulation method from our lab^47^, we overexpressed placental *Igf1* to evaluate its role in placental structure and striatal development. We hypothesized that an overexpression of placental *Igf1* would lead to accelerated growth and maturation of the striatum in a sex-specific manner. We were able to successfully overexpress placental *Igf1* (Igf1-OE) using the aforementioned technique. Interestingly, female and male placentas responded differently to this manipulation, with females showing a promotion of growth and males showing compensatory changes after overexpression with distinct placental structural outcomes. With changes in placenta, Igf1-OE females and males initially had overlapping striatal structure and behavior effects which then diverged in females and males in adulthood. Overall, this study demonstrates that overexpression of placental *Igf1* sex-specifically alters placental structure subsequently altering embryonic striatal development and persistent neurobehavioral changes in mice. This could contribute to the etiology of NDDs, such as ASD and ADHD, in some individuals. This study addresses a significant knowledge gap and expands our understanding of placental functional effects on neurodevelopment. Given the placenta’s greater accessibility than the brain *in utero*, there may be possibilities for leveraging such findings to develop early prevention of pathophysiological mechanisms through placental biology.

## RESULTS

### Placental-targeted CRISPR mediated overexpression resulted in sex-specific phenotypes

After placental-targeted CRISPR manipulation on embryonic day 12 (E12) as previously described in Carver *et al.* 2023,^47^ placentas and offspring were collected on E13, E14, and E18 or allowed to develop postnatally for juvenile and adult behavioral testing prior to adult brain collection (Figure 1A). No change in maternal serum IGF1 protein levels at E14 or E18 was identified in dams that underwent this procedure versus those that underwent sham laparotomies (Supplemental Figure 1A,B). To validate placental *Igf1* overexpression, qPCR analysis was performed for placental *Igf1* expression. At E13, both female and male placentas that received the *Igf1* activation SAM CRISPR plasmid (Igf1-OE) showed an increase in expression in *Igf1* (Figure 1B,C) versus those in the same litters that received the control activation plasmids (Con). At E14, females showed increased placental IGF1 protein levels, while E14 males showed an increase in IGF1 protein in the embryonic body compared to sex-specific controls (Figure 1D-G). This demonstrated an interesting difference in IGF1 protein transport between Igf1-OE female and male samples. Baseline sex differences may influence IGF1 protein transport as control male placentas had more IGF1 protein than control females on E14 (Supplemental Figure 1C). Further evidence of placental *Igf1* overexpression was seen at E14 as Igf1-OE female and male placentas showed a significant or trend decrease of *Igf2*, respectively (Figure 1H,I). Reduced *Igf2* expression in response to excess *Igf1* in the placenta has been previously reported;^5^ this occurs as compensation against redundant function of these two related genes. In sum, this technique was successful in overexpressing placental *Igf1 in vivo,* to be used to determine placental and embryonic developmental consequences. Based on these early sex differences, we evaluated female and male outcomes separately.

**Figure 1.**
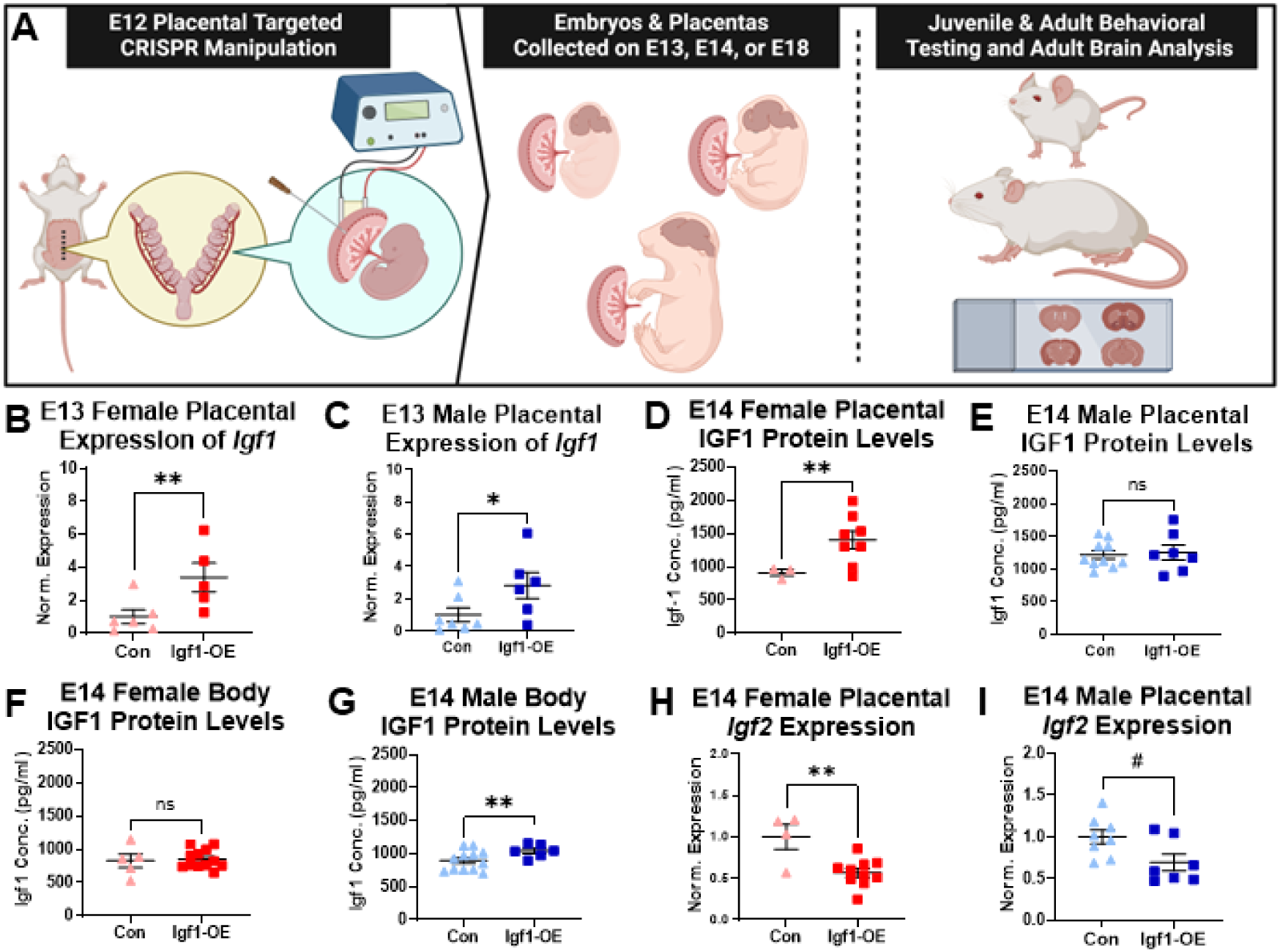
Placental-targeted CRISPR mediated overexpression resulted in sex-specific phenotypes. (A) Schematic of placental-targeted manipulation followed by embryonic and postnatal experiment timepoints. Expression of placental *Igf1* analyzed by qPCR relative to *18s* at E13 in females (B) and males (C) (n=5-7 per group). Placental protein levels of IGF1 by ELISA in E14 females (D) and males (E) (n=3-10 per group). Body levels of IGF1 protein in E14 females (F) and males (G) (n=5-12 per group). Placental *Igf2* expression in females (H) and males (I) (n=4-10 per group). All graphs show mean and SEM. ns=nonsignificant, #p<0.1, *p < 0.05, and **p < 0.01 by linear mixed effects model with litter as a covariate.

### Placental *Igf1* overexpression promoted growth in female placentas

To understand the impact of placental *Igf1* overexpression on the female placenta, analysis of placental gene expression related to angiogenesis and labyrinth growth was analyzed. In the E14 Igf1-OE female placenta, angiogenic factor expression was increased, specifically *Plgf*, *Flt1*, and *Hif-1α* (Figure 2A-C). These three genes are highly associated with angiogenesis, embryonic growth, and placental labyrinth function.^45,48–52^

**Figure 2.**
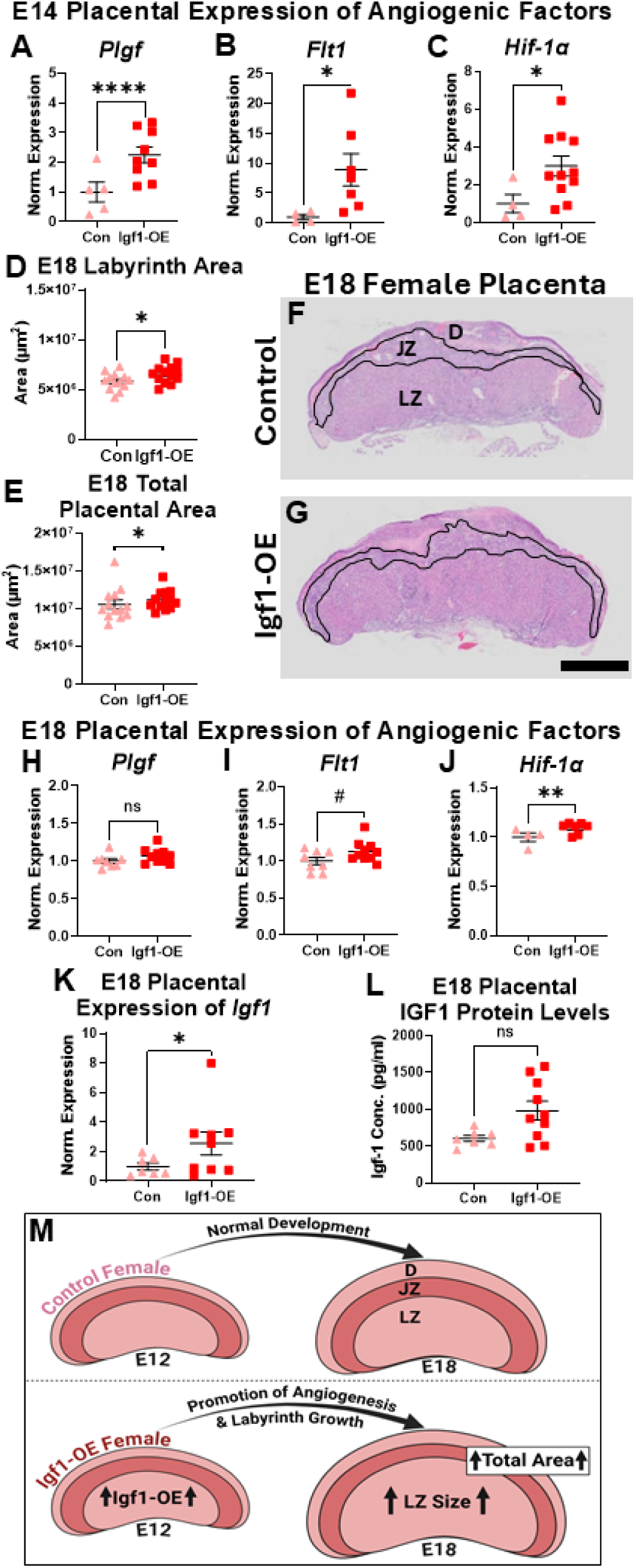
Placental *Igf1* overexpression promoted growth in female placentas. Placental expression of angiogenic factors *Plgf* (A), *Flt1* (B), and *Hif-1α* (C) analyzed by qPCR relative to *18s* in E14 female samples (n=4-11 per group). (D) E18 labyrinth zone area in females (n=13 per group). (E) E18 total placental area in Igf1-OE females (n=13 per group). Representative H&E images of E18 control (F) and Igf1-OE (G) female placental sections. Black line borders the junctional zone, D=decidua, JZ=junctional zone, and LZ=labyrinth zone. E18 placental expression of *Plgf* (H), *Flt1* (I), and *Hif-1α* in female placentas (n=4-10 per group). (K) E18 *Igf1* placental expression (n=7-9 per group). (L) E18 placental IGF1 protein levels (n=7-10 per group). (M) Schematic representing the changes in placental function in Igf1-OE female placentas after placental-targeted *Igf1* manipulation compared to control female placentas. All graphs show mean and SEM. Scale bar represents 1mm. ns=nonsignificant, #p<0.1, *p < 0.05, **p < 0.01, and ****p<0.00001 by linear mixed effects model with litter as a covariate.

Placental structure was also measured from H&E-stained E18 placenta sections. This analysis revealed that labyrinth area and subsequently total placental area were increased in Igf1-OE placentas (Figure 2D-G). Decidua and junctional zone areas were unchanged, indicating a specific effect of placental *Igf1* overexpression on the labyrinth zone in females (Supplemental Figure 2A,B). Further analysis of the E18 placenta showed comparable *Plgf* expression to controls, but a trend increase in *Flt1* expression and a persistent significant increase of *Hif-1α* (Figure 2H-J).

These changes suggested a promotion of placental angiogenesis as well as support for placental and embryonic growth after overexpression of placental *Igf1*. Placental expression of *Igf1* was still higher in E18 females; placental IGF1 protein level at E18 showed change in a similar direction but not at a significant level (Figure 2K,L). There were no changes in placenta mass compared to controls at E13, E14, or E18, suggesting reorganization of placental structure (Supplemental Figure 2C-E). Overall, Igf1-OE female placentas showed that the overexpression of *Igf1* led to a promotion of angiogenesis and labyrinth growth that may facilitate embryonic development (Figure 2M).

### Male placenta downregulated growth after Igf1 overexpression manipulation

Expression of Igf-related genes was analyzed in the male placenta to understand changes after CRISPR manipulation of *Igf1.* In the E14 Igf1-OE male placenta, primary binding protein *Igfbp3*, but not *Igfbp5,* expression was decreased (Figure 3A,B). *Igfbp3* and *Igfbp5* are the two primary binding proteins for IGF1 and increase its bioavailability.^42^ In contrast to this *Igfbp3* decrease in the E14 male placenta, only *Igfbp3* was increased in Igf1-OE females at E18 with no *Igfbp* expression differences otherwise (Supplemental Figure 2F-I). Additionally, expression of *Plgf*, a growth factor involved in labyrinth zone maintenance and embryonic growth,^45,48^ was decreased in E14 Igf1-OE male placentas (Figure 3C). Despite changes in Igf1-OE female placentas, there was no difference in E14 or E18 *Hif-1α* expression or E14 *Flt1* expression in males, but there was a trend decrease in *Flt1* expression at E18 (Supplemental Figure 3A-D). These outcomes indicate early changes to placental signaling that might counteract the initial increase of *Igf1* expression.

**Figure 3.**
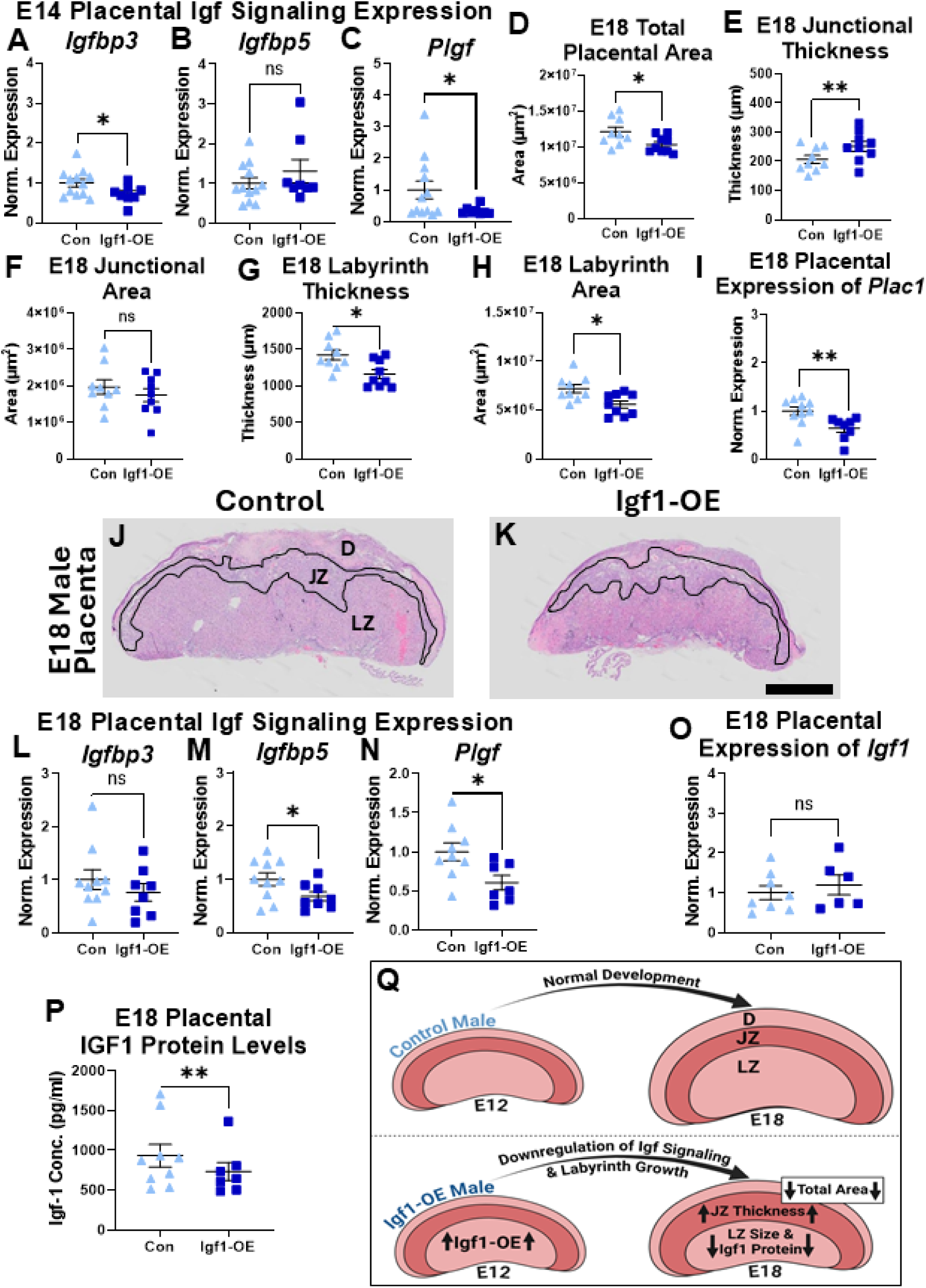
Male placenta downregulated growth after *Igf1* overexpression manipulation. E14 male placental expression by qPCR relative to *18s* for *Igfbp3* (A), *Igbp5* (B), and *Plgf* (C) (n=8-12 per group). (D) E18 total placental area (n=9 per group). (E) E18 junctional zone thickness (n=9 per group). (F) E18 male junctional zone area (n=9 per group). E18 labyrinth thickness (G) and area (H) (n=9 per group). (I) E18 placental *Plac1* expression in males (n=8-10 per group). Representative H&E images of E18 control (J) and Igf1-OE (K) male placental sections. Black line borders the junctional zone, D=decidua, JZ=junctional zone, and LZ=labyrinth zone. E18 male placental expression of *Igfbp3* (L), *Igbp5* (M) and *Plgf* (N) (n=7-10 per group). (O) Placental *Igf1* expression at E18 in males (n=6-8 per group). (P) E18 male placental IGF1 protein levels (n=7-9 per group). (Q) Schematic representing the changes in placental function in Igf1-OE male placentas after placental-targeted *Igf1* manipulation compared to control male placentas. All graphs show mean and SEM. Scale bar represents 1mm. ns=nonsignificant, *p < 0.05 and **p < 0.01 by linear mixed effects model with litter as a covariate.

Later at E18, total placental area was reduced in Igf1-OE males (Figure 3D) due to changes within the subregions. Junctional zone thickness was increased in Igf1-OE males while junctional zone area was unchanged (Figure 3E,F). This demonstrates a change in junctional shape that may alter function. Furthermore, Igf1-OE male placenta labyrinth thickness and area were both decreased (Figure 3G,H). Decidua area was unchanged (Supplemental Figure 3E). Interestingly, E18 *Plac1* expression was also decreased in Igf1-OE male placentas (Figure 3I). A decrease in *Plac1* expression is linked to an expanded junctional zone invading the labyrinth zone, subsequently reducing labyrinth size; decrease in expression is also associated with reduced embryonic body mass.^53^ Representative images of placental structure in E18 control and Igf1-OE males show these structural differences (Figure 3J,K). Despite expression and structural changes in the Igf1-OE male placenta, there was no change in placenta mass compared to controls at E13, E14, or E18, similar to females (Supplemental Figure 3F-H).

Gene expression of downstream IGF signaling factors were assessed at E18 to further understand changes in the placenta that may have contributed to altered structure. In the E18 male placenta *Igfbp3* expression was no longer decreased, but *Igfbp5* was (Figure 3L,M). *Plgf* expression was persistently decreased in E18 Igf1-OE male placentas (Figure 3N). *Igf1* expression was also no longer increased in the E18 Igf1-OE male placenta; in fact, IGF1 protein level in the placenta was decreased (Figure 3O,P). The majority of IGF1 is produced within the labyrinth zone in the placenta;^43,44^ the reduction in this region may have contributed to reduced IGF1 protein production which may reflect compensation for earlier overexpression manipulation (Figure 3Q). The decreased expression of primary *Igfbps* would suggest reduced bioavailability of IGF1 in the placenta, possibly downregulating the IGF signaling pathway. These results indicate that male placentas may have transcriptional and structural changes that maintain *Igf1* within a particular range.

### Placental Igf1-OE led to sex-specific cell cycle changes in the embryonic ganglionic eminence

The observed placental effects were expected to influence embryo development, assessed at E13 and E14. First, body mass was unchanged at E13 in females and males (Supplemental Figure 4A,B). Similarly, ganglionic eminence development, as the primordium of the striatum, was minimally unaffected at E13. Litters collected at E13 were injected with BrdU on E12, approximately 7 hours after CRISPR manipulation of the placenta to capture proliferation changes induced by CRISPR transcriptional activation. Stereology of the E13 ganglionic eminence (Figure 4A) revealed that the rate of cell cycle exit (BrdU+Ki67-/total BrdU+) was unchanged in females but trend increased in Igf1-OE males (Figure 4B,C). Individually, BrdU+ and Ki67+ cell populations were unchanged (Supplemental Figure 4C-F) and the total cell population and volume of the E13 ganglionic eminence were unchanged in females and males (Figure 4D,E and Supplemental Figure 3G,H).

**Figure 4.**
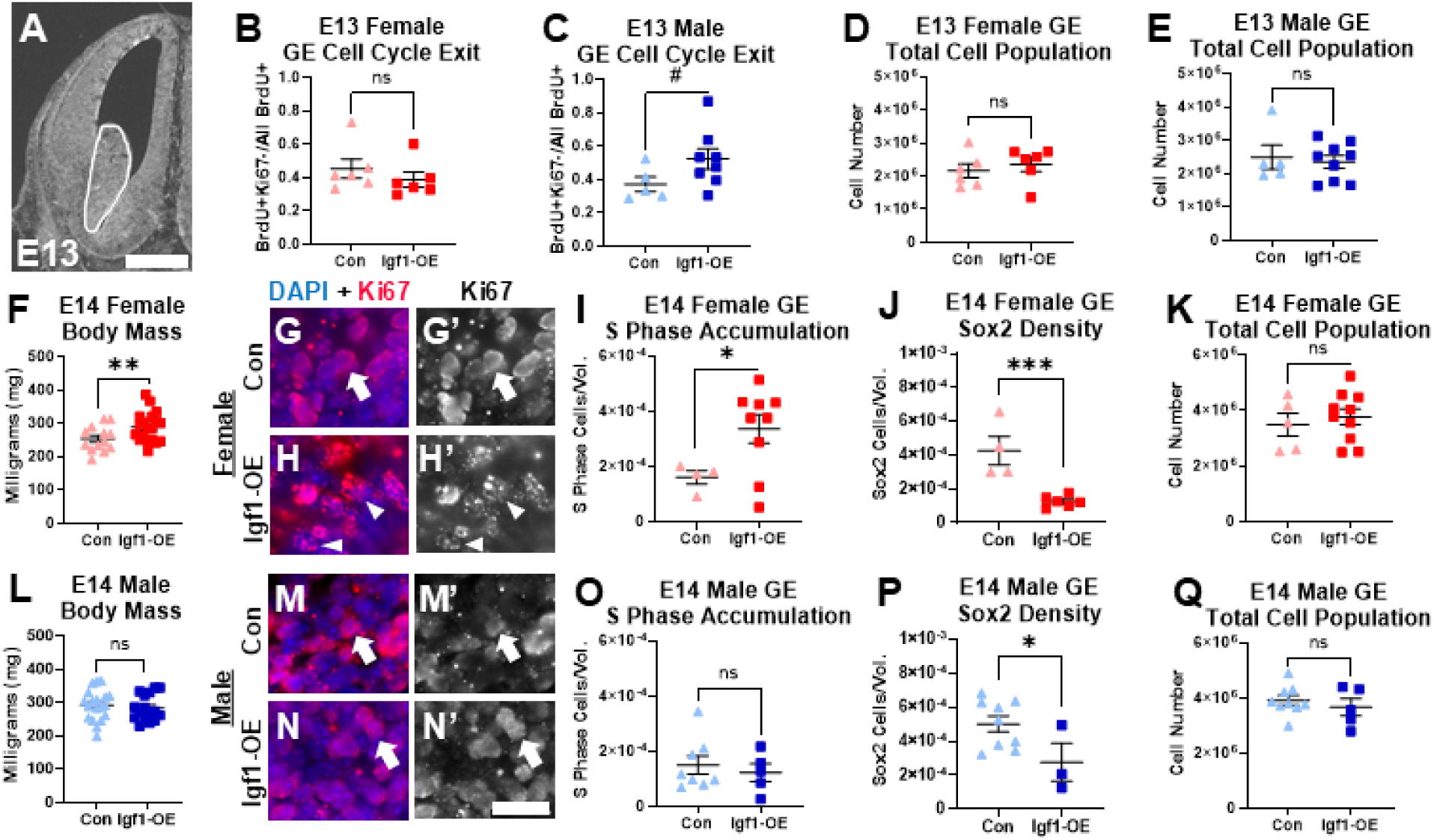
Placental Igf1-OE led to sex-specific cell cycle changes in the embryonic ganglionic eminence. (A) Representative image of a coronal hemi-section of E13 forebrain stained with DAPI; ganglionic eminence (GE) traced in white. Cell cycle exit in the E13 female GE (B) and the E13 male GE (C) (n=5-8 per group). Total cell (DAPI) population in the E13 GE for females (D) and males (E) (n=5-9 per group). (F) E14 female body mass (n=13-19 per group). Representative images of DAPI (blue) and Ki67 (red) staining in the GE: control females (G) and Igf1-OE females (H). The same representative images are shown with Ki67 only in black and white (G’ and H’). White arrows show normal Ki67 diffuse staining and white arrowheads show S phase puncta Ki67 staining. (I) E14 female S phase cell accumulation in the GE (n=4-9 per group). (J) E14 female Sox2+ cell density in the GE (n=4-6 per group). (K) Total cell (DAPI) population in the female GE (n=5-10 per group). (L) E14 male body mass (n=16-19 per group). Representative images of DAPI (blue) and Ki67 (red) staining in the GE: control males (M) and Igf1-OE males (N). The same representative images are shown with Ki67 only in black and white (M’ and N’). (O) S phase cell accumulation in the GE of E14 males (n=5-8 per group). (P) E14 male Sox2+ cell density in the GE (n=3-9 per group). (Q) Total cell (DAPI) population in the male GE (n=5-9 per group). All graphs show mean and SEM. Scale bar represents 500µm in panel A and 20µm in panel N’. ns=nonsignificant, #p<0.1, *p < 0.05, and **p < 0.01 by linear mixed effects model with litter as a covariate.

At E14, body mass was increased in E14 Igf1-OE females but not males (Figure 4F,L), likely influenced by previously described placental changes. Ganglionic eminence was also affected at E14. Ki67+ cell density was assessed, distinguishing between diffuse and punctal staining, representing growth/mitotic and synthesis phases of the cell cycle, respectively.^54^ Ki67+ punctal staining represents DNA replication machinery like other S phase markers, while diffuse staining occurs throughout the cell division cycle.^55–58^ E14 control female ganglionic eminence showed mostly diffuse Ki67 stain (Figure 4G,G’). However, in E14 Igf1-OE females, there was substantial Ki67 punctal staining within the nucleus (Figure 4H,H’). When quantified, Igf1-OE female ganglionic eminence density of S phase cells was higher than controls, suggesting accumulation at this cell cycle stage (Figure 4I). This is consistent with an increased amount of placental IGF1 reaching the embryonic ganglionic eminence since IGF1 is known to promote the G1 to S phase transition in neural precursors.^11^ Sox2+ stem cells in the ganglionic eminence were also decreased in Igf1-OE females (Figure 4J). These findings indicate that the increased IGF1 from placenta advanced ventral forebrain neurogenesis, reducing the progenitor stem cell population. While this might lead to changes in the overall cell population and regional growth, since cell cycle changes were not observed before this time point, total cell population and volume were still unchanged at E14 (Figure 4K and Supplemental Figure 4I). Male embryos also had changes in ganglionic eminence development. Both control and Igf1-OE E14 males had mostly diffuse Ki67+ cells (Figure 4M-N’), and punctal Ki67+ S phase cell density was unchanged (Figure 4O). However, total Sox2+ cell density was also decreased in the ganglionic eminence of E14 Igf1-OE males (Figure 4P), in line with the earlier E13 trend increase in cell cycle exit. Similar to E14 Igf1-OE females, E14 male ganglionic eminence total cell population and volume were unchanged (Figure 4Q and Supplemental Figure 4J). Overall, these results show that cell cycle changes occur in both male and female Igf1-OE ganglionic eminence, with more persistence of the effects in females.

### Placental Igf1-OE affected E18 female and male brain and body growth

Despite placental and ganglionic eminence sex differences, both Igf1-OE females and males had increased striatal volume at E18 (Figure 5A-C). Interestingly, E18 Igf1-OE females had a greater total striatal cell population while Igf1-OE males did not (Figure 5D,E). This difference likely resulted from the different timing of ganglionic eminence cell cycle changes seen in females and males. At this same timepoint, there were no differences in dorsal forebrain volume or cell population across groups (Supplemental Figure 5A-D).

**Figure 5.**
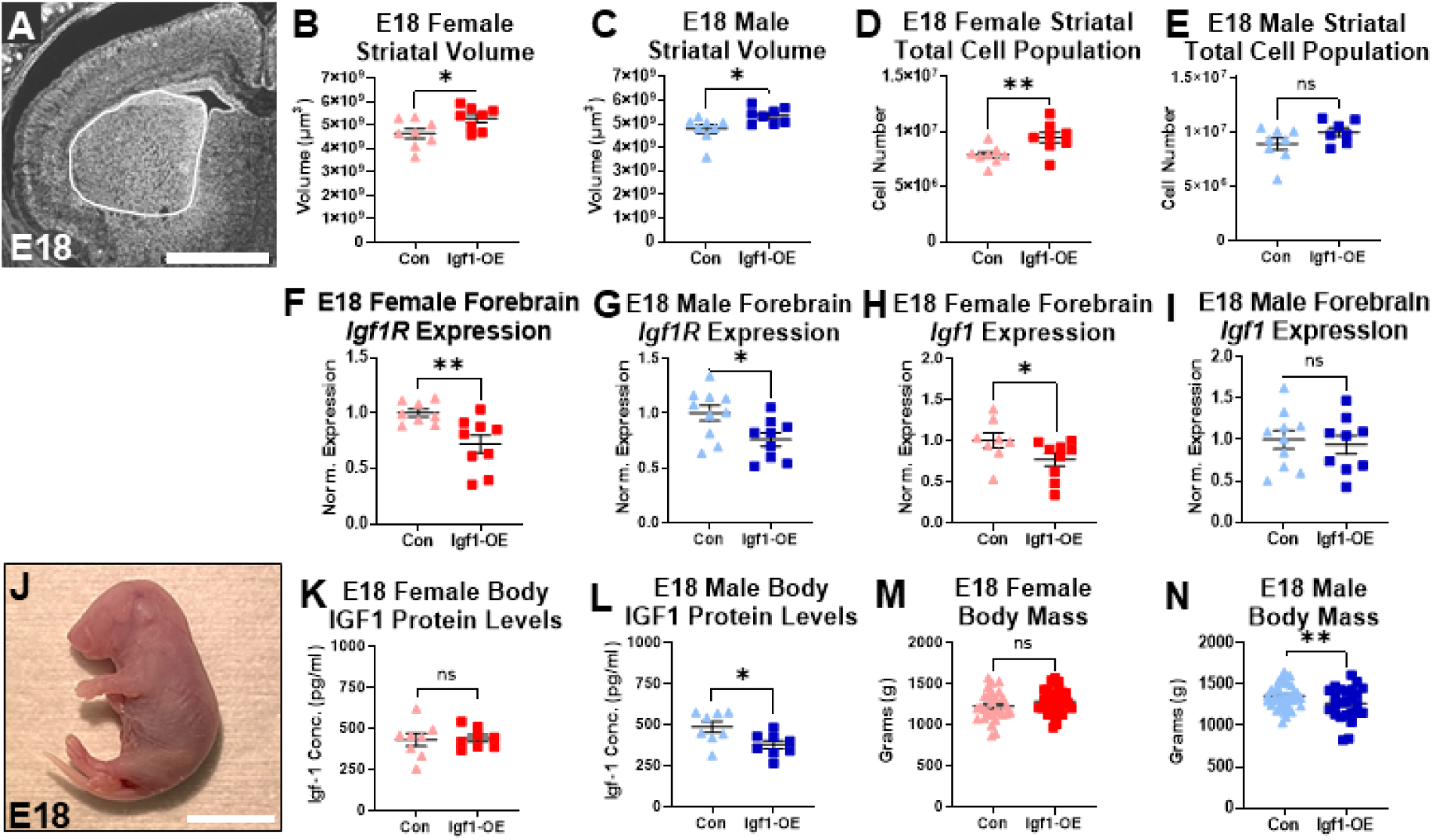
Placental Igf1-OE affected E18 female and male brain and body growth. (A) Representative image of a coronal hemi-section of E18 forebrain stained with DAPI; striatum traced in white. E18 striatal volume for females (B) and males (C) (n=8 per group). Total cell (DAPI) population of the E18 striatum of females (D) and males (E) (n=7-8 per group). E18 forebrain expression of *Igf1R* females (F) and males (G) (n=8-10 per group). E18 forebrain expression of *Igf1* in females (H) and males (I) (n=8-10 per group). (J) Representative image of an E18 embryo body. IGF1 protein levels in the bodies of E18 female embryos (K) and male embryos (L) (n=8 per group). (M) Female body mass at E18 (n=28-35 per group). (N) Male body mass at E18 (n=25-37 per group). All graphs show mean and SEM. Scale bar represents 1mm in panel A and 1cm in panel J. ns=nonsignificant, *p < 0.05 and **p < 0.01 by linear mixed effects model with litter as a covariate.

Manipulated placental *Igf1* overexpression resulted in E18 changes in expression of Igf genes in the brain. Igf1-OE females and males had lower forebrain expression of the primary receptor of IGF1, *Igf1R* (Figure 5F,G). Igf1-OE females, but not males, also had lower *Igf1* in forebrain (Figure 5H, I). This supports a direct response of the brain to placental *Igf1* overexpression. Assessments of E18 body (Figure 5J) however, did not show this impact. E18 IGF1 protein levels in the body (trunk only, head excluded) were unchanged in Igf1-OE female offspring and decreased in Igf1-OE males versus same-sex controls (Figure 5K,L). Similarly, E18 body mass was unchanged in Igf1-OE females and decreased in Igf1-OE males (Figure 5M,N). These results show that altered placental *Igf1* expression may have a particular impact on advancing striatal development as well as influencing general body growth.

### Altered juvenile growth and striatal-dependent behavior in Igf1-OE female and male mice

To examine postnatal outcomes, all placentas within a litter were manipulated with either the Igf1-OE activation plasmid or control plasmid. Since Igf1-OE and control mice were not in the same litter, maternal behavior was evaluated to ensure minimal contributions to later outcomes.^59^ No differences were seen in the maternal behavior of dams that had control or Igf1-OE litters on postnatal day 0 (P0), P3, P8, or P14. (Supplemental Figure 6A-E). Pup body mass also showed no differences at any timepoint prior to weaning on P21 (Supplemental Figure 6F-I). However, both Igf1-OE females and males had increased body length compared to controls of the same sex at P8 and P14 (Supplemental Figure 6J,K and Figure 6A,B) which may reflect the role of placental IGF1 in skeletal development.^60^ Despite altered body length, Igf1-OE pups were not significantly different in early motor development (self-righting) on P6-P10 (Supplemental Figure 6L,M). At P21 and P26, body mass was unchanged in Igf1-OE males, but was larger in Igf1-OE females versus same-sex controls (Supplemental Figure 6N,O and Figure 6C,D).

**Figure 6.**
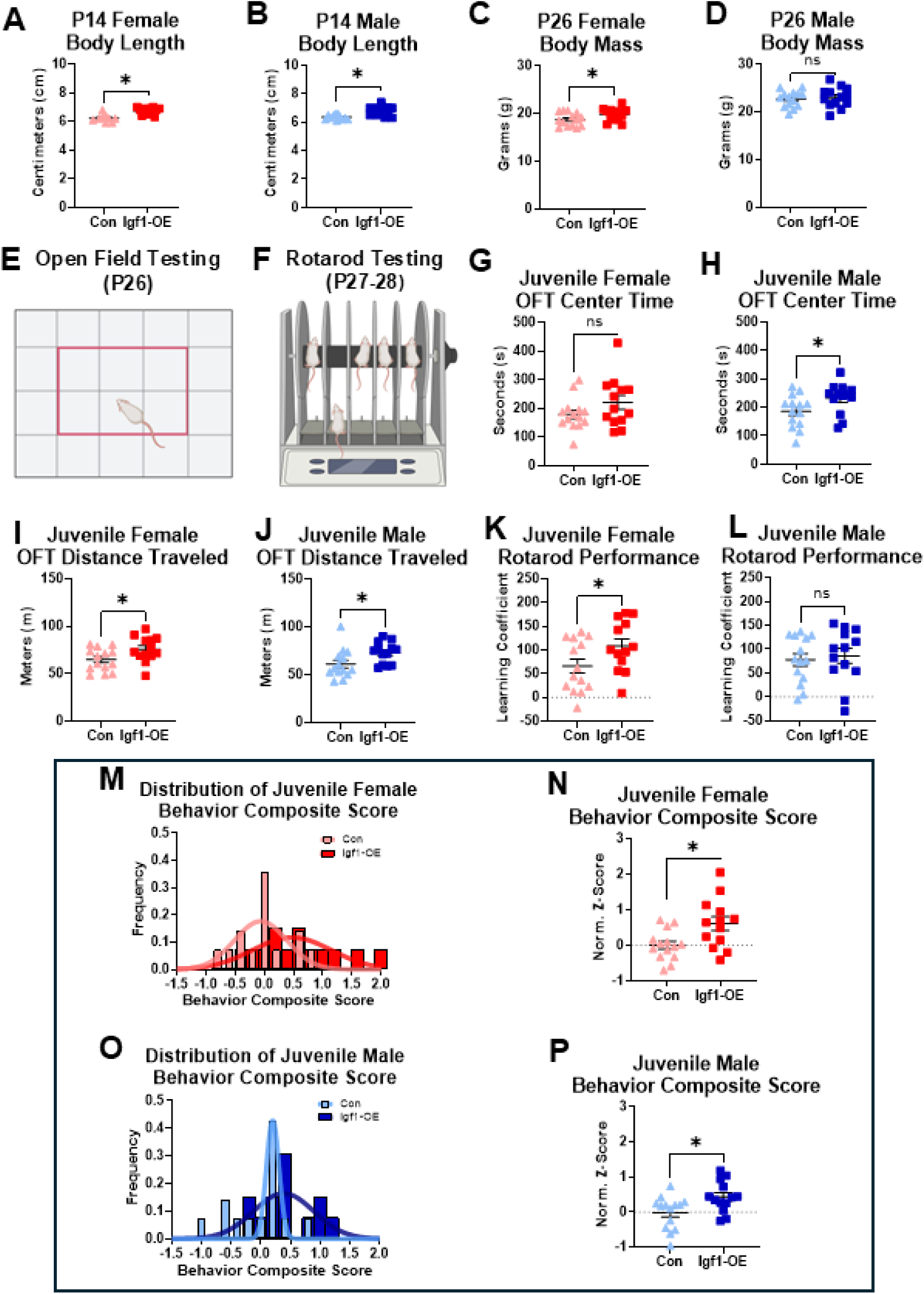
Altered juvenile growth and striatal-dependent behavior in Igf1-OE female and male mice. P14 female (A) and male (B) body length (n=9-15 per group). P26 female (C) and male (D) body mass (n=13-14 per group). Representative images of OFT (E) and rotarod testing (F). (G) Time spent in the center during OFT in P26 females (G) and males (H) (n=13-14 per group). Distance traveled during OFT in P26 females (I) and males (J) (n=13-14 per group). (K) Juvenile female learning coefficient score for rotarod testing (n=13-14 per group). (L) Juvenile male learning coefficient score for rotarod testing (n=13-14 per group). Frequency distribution of juvenile behavior composite score representative of all juvenile behavior tests for females (M) and males (N) versus same sex controls (n=13-14 per group). Juvenile behavior composite scores for females (O) and males (P) (n=13-14). All graphs show mean and SEM. ns=nonsignificant, *p < 0.05 by Welch’s t-test.

Striatal-dependent behavior was evaluated via open field testing (OFT) and rotarod in juvenile mice (Figure 6E,F). On P26, time spent in the OFT center was significantly increased in Igf1-OE males and showed a similar, non-significant effect in females (Figure 6G,H). Both Igf1-OE females and males also showed an increase in distance traveled in the OFT (Figure 6I,J). This may reflect greater impulsive behavior and hyperactive behavior, which are translationally relevant to NDDs.^61,62^ Juvenile mice underwent accelerating rotarod testing P27 to P28, in which they completed 5 trials a day. On the rotarod learning coefficient, Igf1-OE males did not differ from male controls, but Igf1-OE females had increased learning (Figure 6K,L). This Igf1-OE female phenotype is similar to models of NDD risks.^63,64^ When the juvenile behavioral outcomes was combined together as a behavior composite score as done in other studies^30^ for each animal, there was a difference in juvenile behavior in both Igf1-OE females and males versus same-sex controls (Figure 6M-P). These results demonstrate that placental *Igf1* overexpression altered both body growth and striatal outcomes persisting to postnatal juvenile timepoints.

### Persistent changes in adult Igf1-OE female striatal dependent behavior

Adult mice performed behavioral testing beginning at approximately 8 weeks of age (P56). Mice underwent OFT, Y-maze, elevated plus maze (EPM), water T maze, and stereotypy behavioral analysis to test striatal-dependent and other behavior relevant to NDDs (Figure 7A). Adult females and males showed no body mass group differences despite earlier differences in postnatal growth (Supplemental Figure 7A,B). Although juvenile Igf1-OE females and males both were hyperactive and impulsive in the OFT, no group differences were found in adult OFT total distance traveled or center time (Supplemental Figure 7C,D). However, Y maze spontaneous alternation was trend decreased in adult Igf1-OE females (Figure 7B) suggesting poorer working memory than control females. Furthermore, adult Igf1-OE females spent less time in the open arm of the EPM (Figure 7C), exhibiting anxiety-like behavior. Adult Igf1-OE females did not show any difference in trials to reach habit learning or reversal learning criterion in the water T maze (Supplemental Figure 7E-G) but did demonstrate more reversal learning errors in water T maze versus control females (Figure 7D), consistent with reduced flexibility seen in other NDD mouse models.^65^ Adult females had similar levels of stereotypies across groups, including incidences of grooming or total stereotyped behaviors (Figure 7E and Supplemental Figure 7H).

**Figure 7.**
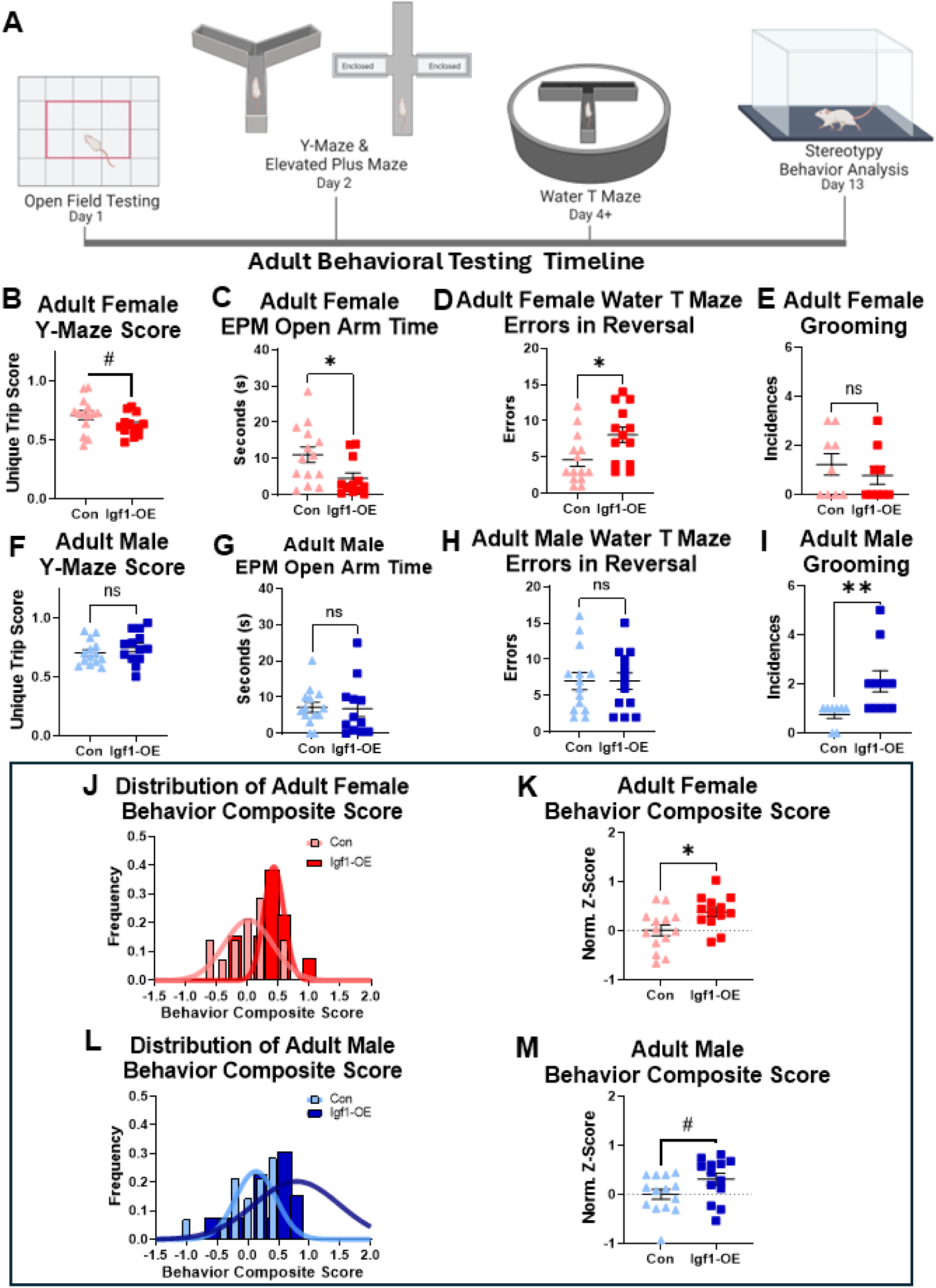
Persistent changes in adult Igf1-OE female striatal dependent behavior. (A) Schematic of adult behavior testing timeline. (B) Adult female Y-maze unique trip score (n=13-14 per group). (C) Adult female time in the open arms during EPM testing (n=12-14 per group). (D) Adult female errors made during water T maze reversal testing (n=13-14 per group). (E) Grooming incidences seen during stereotypy in adult females (n=9 per group). Adult male Y-maze unique trip score (F), time spent in open arm during EPM testing (G), and errors made during water T reversal testing (H) (n=12-14 per group). (I) Adult male grooming incidences during stereotypy testing (n=8-10 per group). (J) Frequency distribution of adult behavior composite score representative of adult behavior tests for Igf1-OE females versus same sex controls (n=13-14 per group). (K) Adult behavior composite scores for females (n=13-14). (L) Frequency distribution of adult behavior composite score representative of adult behavior tests for Igf1-OE males versus same sex controls (n=13-14 per group). (M) Adult behavior composite scores for males (n=13-14). All graphs show mean and SEM. ns=nonsignificant, #p<0.1, *p < 0.05, and **p < 0.01 by Welch’s t-test.

Adult Igf1-OE males had few behavioral differences compared to controls. Igf1-OE males showed no differences in Y-maze spontaneous alternation, EPM open arm time, or water T maze reversal or habit learning (Figure 7F-H and Supplemental Figure 7I-M). At the same time, adult Igf1-OE males had more grooming incidences, and a trend increase in total stereotyped behaviors (Figure 7I and Supplemental Figure 7N). These results indicate a specific increase in repetitive behaviors in adult Igf1-OE males, which has relevance to NDD symptoms.^15,66^

For adults, as for juveniles, the placental-targeted Igf1-OE manipulation altered a behavior composite score generated from all behavioral tests for each animal. Adult Igf1-OE females showed a significant change in behavior composite score compared to control females (Figure 7J,K), and males showed a trend change, driven mostly by increased grooming bouts (Figure 7L,M).

### Altered striatal cell composition only in adult Igf1-OE female brain

Adult brains collected approximately two weeks after the end of adult behavior testing were analyzed for changes in striatal morphology and cell composition (Figure 8A,B). Despite an increase in striatal volume at E18 in females and males, adult striatal volume and total cell population did not differ across groups (Figure 8C-F). The proportion of striatal cells that are neurons was increased in adult Igf1-OE female striatum, a difference not found in males (Figure 8G-H). Additionally, a significant positive correlation in adult Igf1-OE females was found between the neuronal proportion in striatum and the increased reversal learning errors, a behavior associated with striatal function^67^ (Supplemental Figure 8A). These distinctions in females and males may have arisen from the sex differences in early timing of placental functional and ganglionic eminence cell cycle changes. As at earlier timepoints, adult cortical volume and cellular composition of a key subregion, the prefrontal cortex, were also unchanged (Supplemental Figure 8B-I).

**Figure 8.**
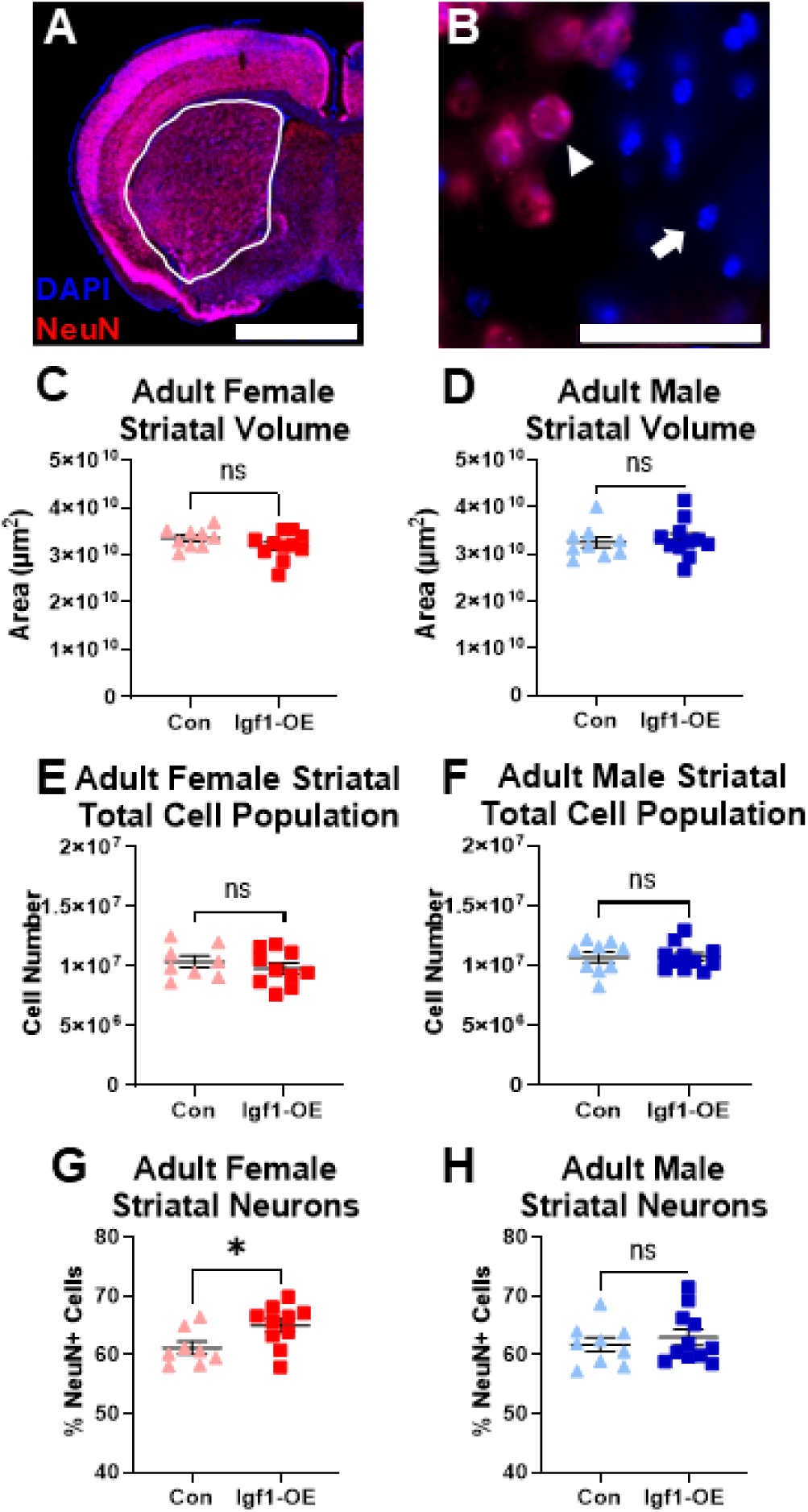
Altered striatal cell composition only in adult Igf1-OE female brain. (A) Representative image of a coronal hemi-section of adult forebrain stained with DAPI (blue) and NeuN (red); striatum traced in white. (B) Representative image of NeuN+ (arrowhead) and NeuN- (arrow) stained cells in the adult striatum. Adult striatal volume for females (C) and males (D) (n=9-11 per group). Total cell (DAPI) population in the adult female (E) and male (F) striatum (n=8-11 per group). Neuron percentage of total cells in the adult female (G) and male (H) striatum (n=8-11 per group). All graphs show mean and SEM. Scale bar represents 2mm in panel A and 50µm in panel B. ns=nonsignificant, *p < 0.05 by Welch’s t-test.

In sum, placental-targeted *Igf1* overexpression altered placental and embryonic brain outcomes differently in females and males resulting in brain and behavioral changes through juvenile and adult life. While there were differences in across females and males, results showed that the effects of altering placental *Igf1* in mice were persistent and relevant to NDD phenotypes.

## DISCUSSION

In this study, we found that placental *Igf1* overexpression altered placental function in a sex-specific manner which influenced striatal development as well as striatal-dependent behaviors emphasizing a strong role for placental biology in long-term brain development. We were able to identify these effects after placental-targeted CRISPR manipulation^47^ that allowed for embryos and postnatal offspring to be studied at multiple timepoints. Through these investigations, we identified responses in female and male placentas that diverged over time after overexpression of endogenous *Igf1*. In both, convergent findings demonstrated that increased placental *Igf1* directly affected embryonic brain likely in conjunction with indirect effects from other placental changes. Additionally, these experiments revealed previously unreported impacts of *Igf1* on striatal development as a result of placental *Igf1* expression, beginning with the embryonic ganglionic eminence. Long-term analysis of these mice further identified changes in juvenile growth and behavior that were similar in females and males. Effects of the placental manipulation on adults differed by sex, with more behavioral and brain impacts in females but some unique striatal-dependent behavioral changes in males. The changes in striatal neurobiology and relevant behaviors are similar to those in mouse models used to study NDDs.^15,68,69^ The findings from this study expand our understanding of placentally produced factors and their role in brain development and potentially as components of NDD pathogenesis. Results of placental *Igf1* overexpression supported the hypothesis that *Igf1* expression advances striatal development, most clearly in female offspring.

First, placental *Igf1* changed as expected in female placenta after CRISPR manipulation; placenta overexpressed *Igf1* within 24 hours, placental IGF1 protein was increased 48 hours later, and overexpression was still evident 6 days later. Placental *Igf1* overexpression was further validated by the expected compensatory decrease in placental *Igf2* by E14. In these mice, overexpression caused a promotion of placental angiogenesis expression and labyrinth zone growth as expected after a promotion of the IGF signaling pathway which regulates these processes (see Figure 2M). Collectively, these placental changes would increase delivery of placental and nutritional factors to the embryo,^70,71^ subsequently altering brain growth, which alongside direct effects of excess IGF1 would also alter striatal development.

Male placentas showed the expected initial increase of *Igf1* expression at E13, but then diverged from expected impacts of the manipulation, particularly with decreased IGF1 protein levels at E18. Reduced expression of primary IGF binding proteins, *Igfbp3* and *Igfbp5*, as well as smaller labyrinth zone size demonstrated likely compensation for early elevated *Igf1*. Decreased *Plgf* at both E14 and E18 may have countered growth promotion from *Igf1* overexpression. The counteraction of growth promotion is further supported by the reduction of *Plac1, a* critical factor for regulating junctional zone invasion within placenta,^53^ concomitant with a junctional zone thickness increase that may have contributed to decreased labyrinth size (see Figure 3Q). Changes to placenta structure also reveal distinct processes of IGF1 protein transport to the embryonic body. IGF binding proteins may influence this, with decreased placental binding protein expression in males leading to IGF1 protein movement out of the placenta.

Why females and males demonstrate these distinct placental responses to CRISPR manipulation is unclear. There are distinct epigenetic factors regulating *Igf1* in placenta that differ by sex which may limit activation by the CRISPR plasmid.^72–74^ Sex differences in placental processes even during normal development are well recognized.^75–77^ For example, there is a higher protein level of placental IGF1 baseline in males versus females which could interact with this manipulation. Additionally, placental levels of IGF1 at E14 and E18 were similar in control males and Igf1-OE females, suggesting a ceiling effect that does not allow for continued overexpression of *Igf1*. At E14, excess IGF1 in Igf1-OE males may be transported out of the placenta, particularly with downregulation of *Igfbp3* and *Igfbp5.* This may account for impacts in males on embryonic striatal volume despite later placental *Igf1* downregulation.

Measurements of body mass and length demonstrate that placental *Igf1* overexpression impacted skeletal development substantially. IGF1 promotes skeletal maturation via bone elongation.^78,79^ Interestingly, Igf1-OE males showed an increase in postnatal skeletal length despite a decrease in body mass observed at E18. The brief placental *Igf1* overexpression observed in males may have altered the trajectory of skeletal development specifically, while placenta compensations against *Igf1* overexpression, such as reduction of *Plgf* and labyrinth zone, may have negatively impacted growth of other parts of the body. When overexpression was consistent throughout gestation, as it was here in females, greater skeletal growth may allow for body mass increase later in development, as seen in juvenile Igf1-OE female offspring.

Direct effect of excess placental IGF1 in female E14 ganglionic eminence was supported by decreased *Igf1* and *Igf1R* brain expression at E18 and S phase cell accumulation in this region at E14. IGF1 promotes the transition of G1 to S phase in embryonic neural progenitors.^11^ Data here is a clear demonstration that placental IGF1 has this impact on the brain *in vivo* and particularly influences development of the primordial striatum. Increased S phase cells at E14 did not imply a total decrease in cell proliferation, as the E18 striatal cell population was enlarged. Other data clarify this finding: excess *Igf1* also reduced Sox2+ stem cells at E14, demonstrating advancement away from a preponderance of self-renewing progenitors to cells fated for neuronal identity. This was supported by adult brain findings in females with a greater proportion of striatal neurons. However, this finding also suggests a reduced glial cell population. Such an impact could result from diminished postnatal gliogenesis in the setting of reduced forebrain *Igf1* and *Igf1R* expression seen at E18. These findings highlight the highly regulated stages of early brain development and show they are intrinsically linked with timing and level of placental growth factor expression. Results from male placenta support this; *Igf1* was only increased early due to sex-specific compensations within the placenta, and postnatal brain had no cell composition changes.

Behavioral differences due to *Igf1* placental overexpression were evident at both juvenile and adult stages. While there were differences in specific functional domains, in general, females showed more extensive alteration compared to same-sex controls, as evidenced by the range of behavior composite scores. This aligns with the more extensive alteration in females of placental *Igf1* expression.

In juveniles, Igf1-OE offspring showed behavior that can be interpreted as more active, more impulsive, and, for females, enhanced rotarod performance, reflecting procedural learning as well as more tendency to repetitive behavior. These effects did not persist into adulthood, approximately four weeks later. With the clear influence of Igf1-OE on embryonic striatal growth, striatal enlargement may have contributed to behavioral abnormalities, as shown in some models of NDD risk.^15,68^ In some individuals, increased striatal volume is associated with increased restrictive, repetitive behaviors, similar to the findings of this study.^80^ One possible contributor to these phenotypes appearing only in juveniles may have been the shorter time elapsed since placental effects. Furthermore, all animals underwent both juvenile and adult testing which likely influenced the age-dependent differences, especially in OFT, as the novelty of the task may have contributed at the juvenile timepoint. In addition, changes in body growth may also contribute, however body length and mass were not correlated with juvenile behavioral outcomes that are highly dependent on motor development (data not shown). It is also interesting to note that juvenile behavioral deficits that improve in adulthood is a phenomenon observed in some individuals with NDDs.^81^

In adulthood, behaviors affected by placental Igf1-OE were distinct in females and males, as were brain outcomes. Males, which had no clear striatal volume or cellular abnormalities, had increased stereotyped behaviors. Multiple neuronal and glial alterations have been implicated in these phenotypes,^82–84^ some of which may be influenced by developmental changes in *Igf1*. Females, which had disproportionately more striatal neurons and larger prefrontal cortex, demonstrated deficits in the reversal phase of striatal-dependent water T-maze. Females also showed increased anxiety-like behavior and reduced working memory which are not often associated with changes to the striatum but are often found in models of NDDs. Mouse models of NDDs have linked striatal and prefrontal abnormalities with similar behavioral findings.^69,85–87^ These outcomes may be indicative of other brain changes, since the overexpression of placental *Igf1* likely had broad effects. Overall, changes in postnatal brain and behavior demonstrate a link between placental *Igf1* expression and the development of NDD relevant phenotypes.

NDDs are extremely common developmental conditions, including ASD which affects about one in every 36 children^88^ and ADHD which affects 11.3% of children.^89^ NDDs such as ASD and ADHD have complex etiologies influenced by environmental-genetic interactions, including the *in utero* environment in which placental function is particularly important.^31,90–92^ They also involve developmental impacts on the striatum and striatal-dependent behavior.^15,22,23^ The prevalence of NDDs has increased as seen by an approximately 4% increase in children diagnosed with ADHD over a 20-year period^93^ and a staggering 175% increase in ASD diagnoses from 2011 to 2022.^94^ The rapidly increasing prevalence of NDDs further necessitates research on contributing mechanisms including those from placental biology. This study demonstrates that placental *Igf1* expression can alter striatal development and striatal-dependent behavior postnatally. The changes observed, such as increased striatal volume, hyperactivity, and altered learning and memory are relevant to NDDs.^15^ This shows that placental *Igf1* is highly influential for neurodevelopment; changes to expression of this essential factor may have a role in NDD etiology in people. Furthermore, this study focuses on sex-differences in this model that revealed distinct sex-specific phenotypes highlighting an important factor that must be considered in neuroplacentology studies. Greater understanding of placenta to brain mechanisms is just as critical as studies that focus on brain development in isolation. We were able to perform this experiment by using *in vivo* placental-targeted CRISPR manipulation^47^ which allowed for the first study of placental specific *Igf1* overexpression and its impact on neurodevelopment in mice. This technique was ideal for a study of this kind as it demonstrates how the placenta can be manipulated to alter the trajectory of striatal development and function, but this technique could also be used to study changes to other tissues, such as heart as the placenta-brain-heart axis is an emerging field of study.^95,96^ The findings from this study will be an important part of current medical investigations into placental factors that may be applied to individuals with perinatal risks. Just one example is the delivery of IGF1 using nanoparticles to the placenta in non-human primates to address intrauterine growth restriction.^97^ This study and others demonstrate a necessity for more research of this kind as well as mechanisms for developing preventive interventions; placental-targeted interventions are more feasible and less invasive than directly manipulating the developing fetal brain. Our study will aid in the understanding of placental *Igf1* and its implications for lifelong trajectories particularly relevant to NDDs.

### Limitations of the study

Placental CRISPR manipulation contributes to understanding placental produced IGF1’s impact on brain development, but this manipulation does not mimic a “usual” environment as expressed levels may be outside normal physiological ranges, in cells which may not typically express it, and the procedure used for CRISPR introduction was invasive. Despite this, we were able to garner a better understanding of placental *Igf1* expression on fetal brain development and we did not notice poor health in embryos^47^ or postnatal animals that underwent this manipulation. Additionally, brain regional structures and cellular populations were not comprehensively assessed as this study focused on striatal outcomes. Furthermore, male placenta was less responsive to CRISPR manipulation for unclear reasons. This may be specific to *Igf1* manipulation; a similar study in which placental *Plgf* was overexpressed did not report sex-specific expression differences.^98^ Lastly, placental *Igf1* expression was manipulated in mid-gestation (E12), so this study does is not representative of earlier changes to placental *Igf1* expression.

## RESOURCE AVAILABILITY

### Lead contact

Further information and requests for resources and reagents should be directed to and will be fulfilled by the lead contact, Dr. Hanna E. Stevens.

### Materials availability

This study did not generate new unique reagents.

### Data and code availability

Any additional information required to reanalyze the data reported in this paper is available from the lead contact upon request.

## ACKNOWLEDGMENTS

The authors acknowledge the following funding sources: R01 MH122435 (H.E.S), NIH T32GM008629 (A.J.C.), and NIH T32GM145441 (A.J.C.) as well as the University of Iowa Graduate College (A.J.C), Biomedical Summer Undergraduate Research Program (F.M.F.) and the Beginning and Early Stage Translational Research Program (A.G.). The authors would like to thank Dr. Benjamin Hing for assistance with statistics. The authors would also like to thank Dr. Val Sheffield and Dr. Calvin Carter’s labs at the University of Iowa for the use of their surgery room and equipment. The graphical abstract, figure 1A, figure 2M, figure 3Q, figure 6E, figure 6F, and figure 7A were created with BioRender.com Carver, A. (2025) https://BioRender.com/y9rz1lx, https://BioRender.com/cjwnahr, https://BioRender.com/dbxv36w, https://BioRender.com/vqoypfh, and https://BioRender.com/pb6ok5n.

## AUTHOR CONTRIBUTIONS

Conceptualization, A.J.C. and H.E.S.; data curation, A.J.C.; formal analysis, A.J.C. and H.E.S.; funding acquisition A.J.C., F.M.F., A.G., H.E.S.; investigation A.J.C., F.M.F., R.J.T., S.B., N.R.K., and A.G.; methodology, A.J.C., R.J.T., and H.E.S.; project administration, A.J.C., R.J.T., and H.E.S.; resources, H.E.S.; supervision, A.J.C. and H.E.S.; validation, A.J.C., F.M.F., and R.J.T.; visualization, A.J.C., F.M.F., S.G., N.R.K., and A.G.; writing—original draft, A.J.C., F.M.F., R.J.T., S.B., N.R.K., and A.G.; writing—review & editing, A.J.C., F.M.F., R.J.T., and H.E.S.

## DECLARATION OF INTERESTS

The authors have no interests to declare.

## DECLARATION OF GENERATIVE AI AND AI-ASSISTED TECHNOLOGIES

None.

## STAR★METHODS

### KEY RESOURCES TABLE

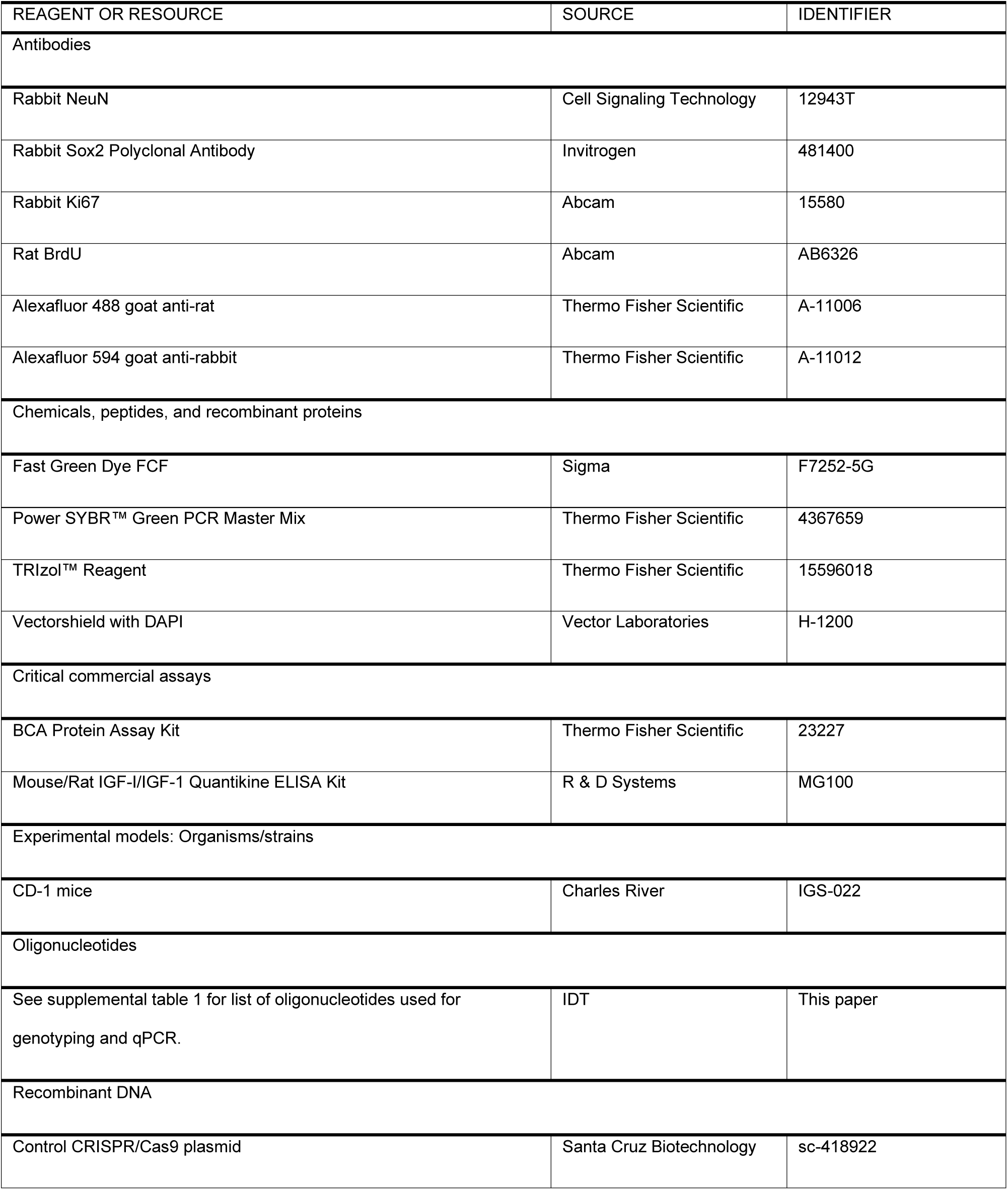
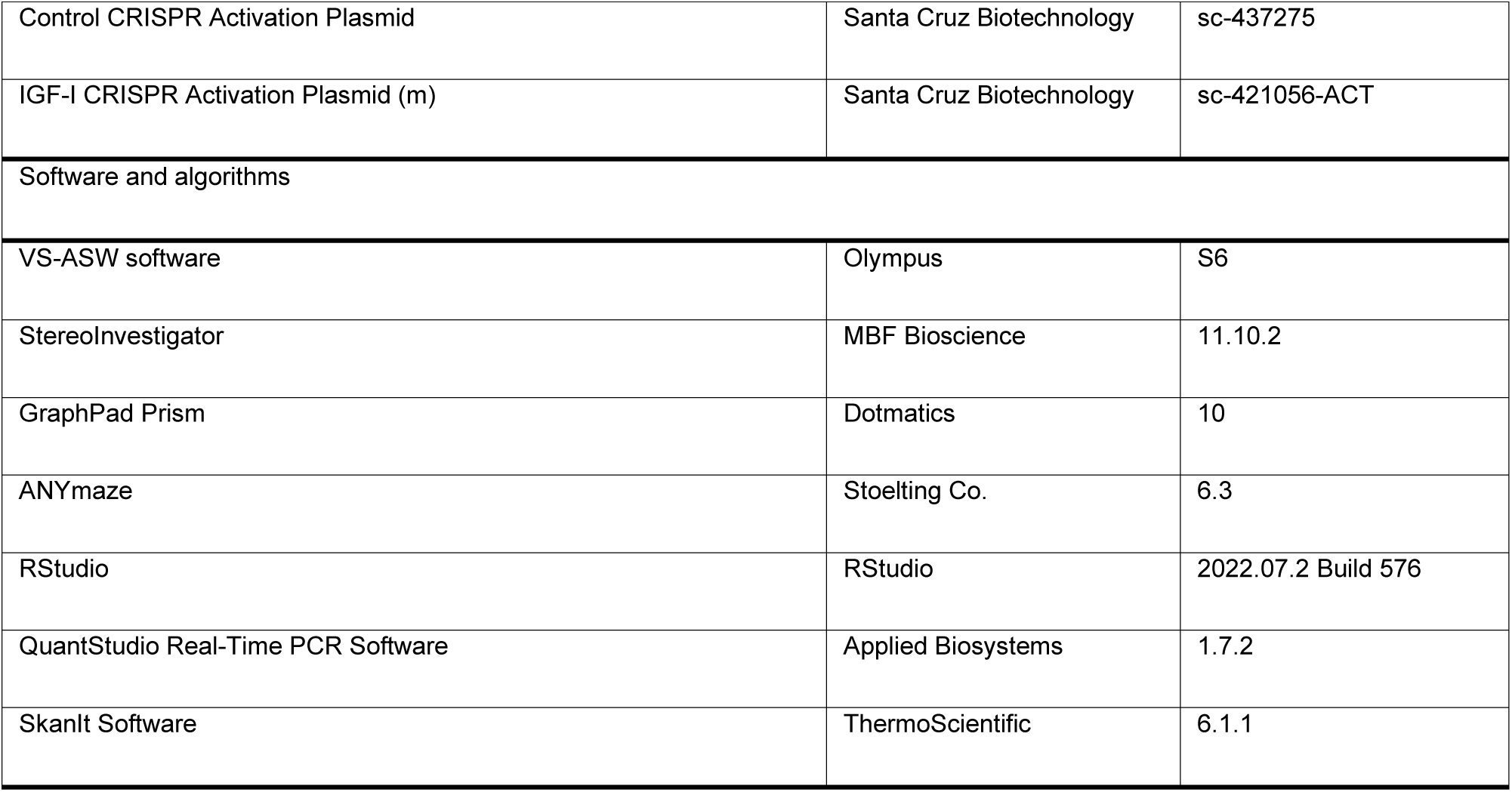

### EXPERIMENTAL MODEL DETAILS

CD-1 mice were bred in-house or ordered from Charles River. 8–16 week-old females were bred to CD-1 males and the identification of a copulation plug was used to define E0. After identification of a plug, the dam was singly housed. Embryos and placentas were collected on E13, E14, and E18 (See below for details). Offspring were weaned and group-housed at P21, monitored P0-28, and underwent juvenile behavior testing P26-28. Offspring mice also underwent adult behavioral testing between 8 and 12 weeks of age. Offspring mice underwent euthanasia, perfusion, and collection between 12 to 14 weeks of age. Both female and male offspring were used for all embryonic and postnatal assessments. Mouse care, handling, husbandry, and assessment were performed as approved by University of Iowa IACUC. Mice were housed in a 12-hour light cycle with food and water ad libitum.

### METHOD DETAILS

#### Placental Targeted CRISPR Manipulation for Embryonic Collection

Placental targeted CRISPR manipulation was performed as described in Carver *et al.* 2023.^47^ On E12, CD-1 dams were anesthetized under isoflurane. A laparotomy was performed to expose the uterine horns, then three pairs of adjacent placentas spaced across the uterine horns were chosen for manipulation. Using a glass microneedle, one placenta in each pair was injected with a control CRISPR plasmid (Control CRISPR/Cas9 Plasmid only in some E14 collections otherwise Control CRISPR Activation Plasmid, Santa Cruz Biotechnology for all time points) and the other was injected with IGF-I CRISPR Activation Plasmid (m) Santa Cruz Biotechnology). No difference between these two CRISPR control plasmids was identified, as reported in Carver *et al.* 2023.^47^ Placentas were injected with plasmids then electroporated. Manipulated placenta locations were recorded so that they could be identified during embryonic collection. Post-procedure, the dam was allowed to recover until embryonic collection. Litters that were collected on E13 received a 10 mg/kg BrdU injection approximately 7 hours post procedure. Placentas and embryos were collected and weighed at E13, E14, or E18 after dam euthanasia.

#### Sex Genotyping

PCR with either *Jarid* or *Rmb31* primers (Supplemental table 1) was used to determine sex.

#### Quantitative Polymerase Chain Reaction

E13, E14, and E18 placentas were collected, minced into smaller pieces, and stored in RNAlater at - 80°C. Pieces from across the placenta were used for each assay to get more representative expression of placental genes as proximity to CRISPR injection site could have led to variation in expression across the placenta. At E18 forebrain was isolated, placed into RNAlater, and stored at −80°C. RNA was isolated from placenta and forebrain via Trizol method. Total RNA was normalized prior to cDNA synthesis. qPCR was conducted with PowerSYBR Master Mix on Thermo Fisher Scientific ViiA 7 Real-Time PCR System (Thermo Scientific). *18s* was used as the housekeeping gene for placentas while *Gapdh* was used as the housekeeping gene for forebrains. See supplemental table 1 for primer sequences. Genes were normalized to the appropriate housekeeping gene and analyzed using the ddCT method to calculate normalized fold change relative to controls.

#### ELISA Protein Analysis

After collection at E14 and E18, placentas were minced as described for qPCR and stored at −80°C. At E14 and E18 bodies (trunk only, head removed) were also stored at −80°C. E14 and E18 placentas and E14 bodies were homogenized as previously described in Carver *et al.* 2023.^47^ E18 bodies were homogenized via bead homogenization. Total protein in the placentas and bodies were quantified using BCA Protein Assay Kit (Thermo Fisher Scientific, 23227) following the manufacturer instructions. Samples were then homogenized to the same total protein concentration and then were analyzed in duplicate using the Mouse/Rat IGF-I/IGF-1 Quantikine ELISA Kit (R&D MG100) following the manufacturer instructions. Absorbance measures were recorded for ELISA plates on a plate reader and results were analyzed on the SkanIt Software (Thermo Scientific).

#### Placental Histology

E18 placentas for histological analysis were cut down the midline; half was placed into 10% neutral buffered formalin and stored at 4°C until paraffin embedded. Placentas were vibratome sectioned by the University of Iowa Comparative Pathology and Histology Research Core at 4 µm from the midline cut so that all three regions were visible. Slides were H&E stained and mounted with Cytoseal Mounting Media. Placental sections were visualized via bright-field microscopy on Axio Imager M.2 microscope (Zeiss). The labyrinth zone, junctional zone, and decidua were measured in two adjacent sections from each sample using StereoInvestigator (Microbrightfield) and averaged. All placental histology measures were performed by the same blinded experimenter.

#### Embryonic Brain Analysis

E13, E14, and E18 heads were fixed in 4% PFA for up to 72 hours at 4°C, cryoprotected in 20% sucrose, and frozen in O.C.T. and stored at −80°C until sectioned. Embryonic heads were serial sectioned coronally at 25 µm onto slides and then immunostained with primary antibodies for Ki67 (1:200), Sox2 (1:200) and BrdU (1:500) (after acid incubation for BrdU only) and appropriate secondary antibodies (1:500). Slides were mounted with VectaShield with DAPI (Vector Labs). On an Axio Imager M.2 microscope (Zeiss), striatum, cortex, and forebrain were assessed with unbiased stereology (StereoInvestigator) to estimate volume, cell population, and density measures for Ki67+, BrdU+, Sox2+, and DAPI+ cells.

#### Placental Targeted CRISPR Manipulation for Postnatal Study

For postnatal evaluation of placentally manipulated litters, the placental targeted CRISPR manipulation was performed as described above except all placentas in the litter received one type of plasmid. All placentas were then electroporated and maternal dams were allowed to recover as previously described. Maternal dams were monitored daily and allowed to carry litter to term.

#### Analysis of Maternal Behavior

Dams that gave birth to manipulated litters were monitored for maternal care behaviors. On P0, P3, P8, and P14, dams were recorded for 24, 3, 2, and 1 hours respectively. Recordings at P3, P8, and P14 took place only during the light cycle. Videos were analyzed in four-minute increments and scored by the same blinded experimenter for time on or off pups as previously described in Schroeder *et al.* 2022.^59^

#### Juvenile Postnatal Measures

After maternal behavior recording, pup weights and lengths (nose to rump) were recorded on P8 and P14.

#### Postnatal Self-Righting Motor Test

Self-righting testing was performed daily from P6 to P10 for pups that underwent placental manipulation. Pups were removed from the home cage and then gently laid on their back and the time to return to their feet was recorded. If the task was completed in under a second, the result was recorded as one second.

#### Behavioral Testing

A maximum of five mice of one sex from each litter were used for behavior after weaning. Prior to each behavioral test, mice were allowed to habituate to the testing location in their home cage for a minimum of 30 minutes. All juvenile (starting at P26) and adult (starting at P56-70) offspring behavioral testing was performed during the light cycle.

#### Open Field Testing

Individual juvenile and adult mice were placed into a plexiglass open field testing apparatus and recorded overhead for 30 minutes using ANYMaze (Stoelting) to monitor behaviors on a single day. After testing, mice were returned to their home cage with their same sex littermates.

#### Rotarod Testing

Mice were tested on a rotarod apparatus (Ugo Basile) for five trials a day for two days. The rotarod apparatus was set to accelerate from 4 to 80 rpm over 240 seconds. The end of a trial was determined when the mouse either fell off the rotarod or after two full revolutions. A learning coefficient was calculated by subtracting the average time of the last two trials (trials 4 and 5 from day two) by the average time of the first 2 trials (trials 1 and 2 from day one).

#### Y-Maze Testing

Mice were placed into a plexiglass Y-maze apparatus and monitored for 5 minutes. All mice were placed into the same location within the Y-maze apparatus at the beginning of testing to reduce variability in results. Entry into each arm was recorded and a spontaneous alternation score calculated as the proportion of unique arm entry triplets to total arm entry triplets.

#### Elevated Plus Maze Testing

Mice were placed into the center of the EPM apparatus and monitored for 5 minutes using ANYMaze to monitor behavior including time spent in open arm zones.

#### Water T Maze Testing

Water T maze testing was performed in a room with distal visual cues and a tub with room temperature water dyed white with non-toxic paint to obscure the view of the white platform placed at the end of one of the arms of the T maze as previously described in Maurer *et al.* 2025.^85^ Mice first underwent training to swim from the base of the T maze to the platform in either the left or right arm (counterbalanced); time and errors (swimming into the wrong arm) were assessed on up to 10 trials a day. If mice took longer than 60 seconds to identify the platform, the trial was terminated. Once mice found the platform five trials in a row without error in one day (criterion), training continues for two more days. Reversal testing was then performed by switching the position of the platform to the opposite arm. Time to reach the platform was then measured and errors recorded across 10 trials a day until the same criterion was reached again. The number of trials to training and reversal criterion were corrected by subtracting the first four trials of each day until criterion was met.

#### Stereotyped Behavior Analysis

Mice were placed into an open field plexiglass testing box with one transparent wall and video recorded from the side for 20 minutes. Numbers of behaviors were scored by the same blinded experimenter. For one minute in each 5-minute interval, including. grooming bouts, rearing, cage circling, and self-circling. A total stereotyped behavior score was calculated from normalized z-scores of each individual behavior. Individual behavior counts and the total score were used for statistical analysis.

#### Adult Brain Analysis

Adult brains were collected at least two weeks after the end of behavioral testing using transcardial perfusion with saline and then 4% PFA. Brains were dissected out, postfixed in 4% PFA for approximately 3 days, and then cryoprotected in 20% sucrose in PBS at 4°C until they could be embedded in O.C.T. and stored at −80°C. Brains were serially sectioned coronally at 50 µm, and then immunostained with NeuN primary antibody (1:300) and the appropriate secondary antibody (1:500). Stained sections were mounted onto slides with VectaShield with DAPI. On an Axio Imager M.2 microscope coupled to StereoInvestigator, striatum, cortex, and prefrontal cortex were assessed and unbiased stereology was performed to measure cell populations and densities for NeuN+ and DAPI+ cells. by the same blinded experimenter.

### QUANTIFICATION AND STATISTICAL ANALYSIS

All offspring outcomes were analyzed separately by sex, run through the ROUT outlier test in GraphPad (Prism), and statistical outliers were excluded prior to statistical comparisons. Outlier tests were not performed for maternal behavior.

#### Embryonic and Placental Analyses

Embryonic and placental analyses were run through the linear mixed effects model (lmer function) in R to incorporate litter as a covariate. The two comparisons of E14 female placental IGF1 levels and E14 male versus female control placental IGF1 levels were not compared using the lmer function in R as sample size was too small, so an unpaired Welch t-test in GraphPad was used for these two comparisons.

#### Postnatal and Maternal Analyses

Postnatal group comparisons were made using Welch t-tests in GraphPad. Maternal results were analyzed using an unpaired t-test in GraphPad. Correlations were performed as a simple linear regression in GraphPad.

#### Behavior Composite Score

Behavior composite score was calculated for each animal by generating a z-score for each measure and then averaging individual z-scores across measures for juvenile or adult time points, as described in Vacher *et al.* 2021.^30^ Absolute values of z-scores were used to adjust negative average z-score values to focus on degree of differences compared to controls. The juvenile behavior composite score was calculated as an average of the z-score of total open field center time, total open field distance traveled, and rotarod learning coefficient. The adult behavior composite score was calculated as an average of the z-score of total open field center time, total open field distance traveled, Y-maze spontaneous alternation, EPM open arm time, errors made in water T maze training, errors made in water T maze reversal, and total stereotyped behaviors from stereotypy analysis.

### ADDITIONAL RESOURCES

To better understand the placental-targeted CRISPR manipulation used in this paper see the attached link to the published protocol paper and informational video: https://app.jove.com/t/64760/mouse-in-vivo-placental-targeted-crispr-manipulation

## Notes

### Competing Interest Statement

The authors have declared no competing interest.

