## Supplemental Material for "Placental *Igf1* Overexpression Sex-Specifically Impacts Mouse Placenta Structure, Altering Offspring Striatal Development and Behavior"

#### SUPPLEMENTAL TABLE 1

| | |
|---|---|
| ***Gapdh* Forward** | 5’-GGTGAAGGTCGGTGTGAACG-3’ |
| ***Gapdh* Reverse** | 5’-CTCGCTCCTGGAAGATGGTG-3’ |
| ***Igf1* Forward** | 5’-ATTGCTCTAACATCTCCCATCTCT-3’ |
| ***Igf1* Reverse** | 5’-GAAATGAATTGGTGGGCAGGG-3’ |
| ***Igfbp3* Forward** | 5’-CCAGGAAACATCAGTGAGTCC-3’ |
| ***Igfbp3* Reverse** | 5’-GGATGGAACTTGGAATCGGTCA-3’ |
| ***Igfbp5* Forward** | 5’-AAGACAGCACAACTTCAGTTCA-3’ |
| ***Igfbp5* Reverse** | 5’-TCACAGGGAAGGATGTTTGAGT-3’ |
| ***Plgf* Forward** | 5’-TGAAGGCATGTAGAGGGGAC-3’ |
| ***Plgf* Reverse** | 5’-CACTCTGCCTGTGTTCCAGA-3’ |
| ***Hif-1α* Forward** | 5’-AGCCCTAGATGGCTTTGTGA-3’ |
| ***Hif-1α* Reverse** | 5’-TATCGAGGCTGTGTCGACTG-3’ |
| ***Flt1* Forward** | 5’-CCTCCGTGCATGTGTATGAA-3’ |
| ***Flt1* Reverse** | 5’-CATCCTCGGTTGTCACATCTT-3’ |
| ***Igf1R* Forward** | 5’-GTGGGGGCTCGTGTTTCTC-3’ |
| ***Igf1R* Reverse** | 5’-GATCACCGTGCAGTTTTCCA-3’ |
| ***Jarid 1C*** | 5’-CTGAAGCCTTTGGCTTTGAG-3’ |
| ***Jarid 1D*** | 5’-CCACTGCCAAATTCTTTGG-3’ |
| ***Rbm31* Forward** | 5’-CACCTTAAGAACAAGCCAATACA-3’ |
| ***Rbm31* Reverse** | 5’-GGCTTGTCCTGAAAACATTTGG-3’ |

**Supplemental Table 1: Oligonucleotide/Primer information.** List of oligonucleotide/primer sequences used in this paper for genotyping or qPCR.

#### SUPPLEMENTAL FIGURE 1

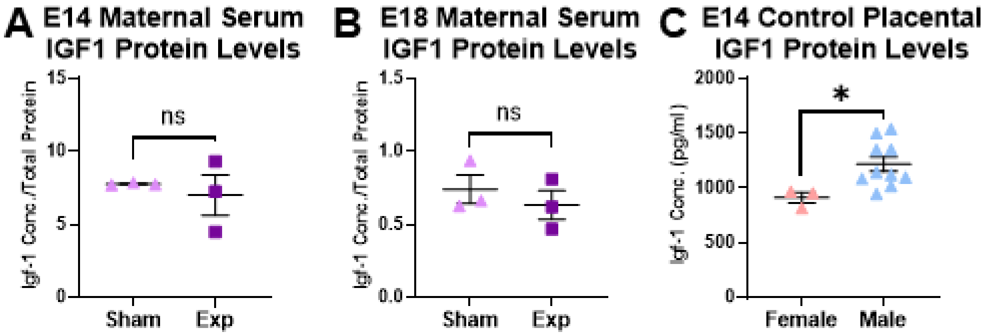


**Supplemental Figure 1: IGF1 protein analyses.** IGF1 protein levels in serum at E14 (A) and E18 (B) in pregnant dams that either underwent sham laparotomies (sham) or placental-targeted CRISPR manipulation (Exp) (n=3 per group). IGF1 protein levels in E14 female control placentas compared to E14 control male placentas (n=3-10 per group). All graphs show mean and SEM. ns=nonsignificant, *$p < 0.05$ by Welch's t-test.

#### SUPPLEMENTAL FIGURE 2

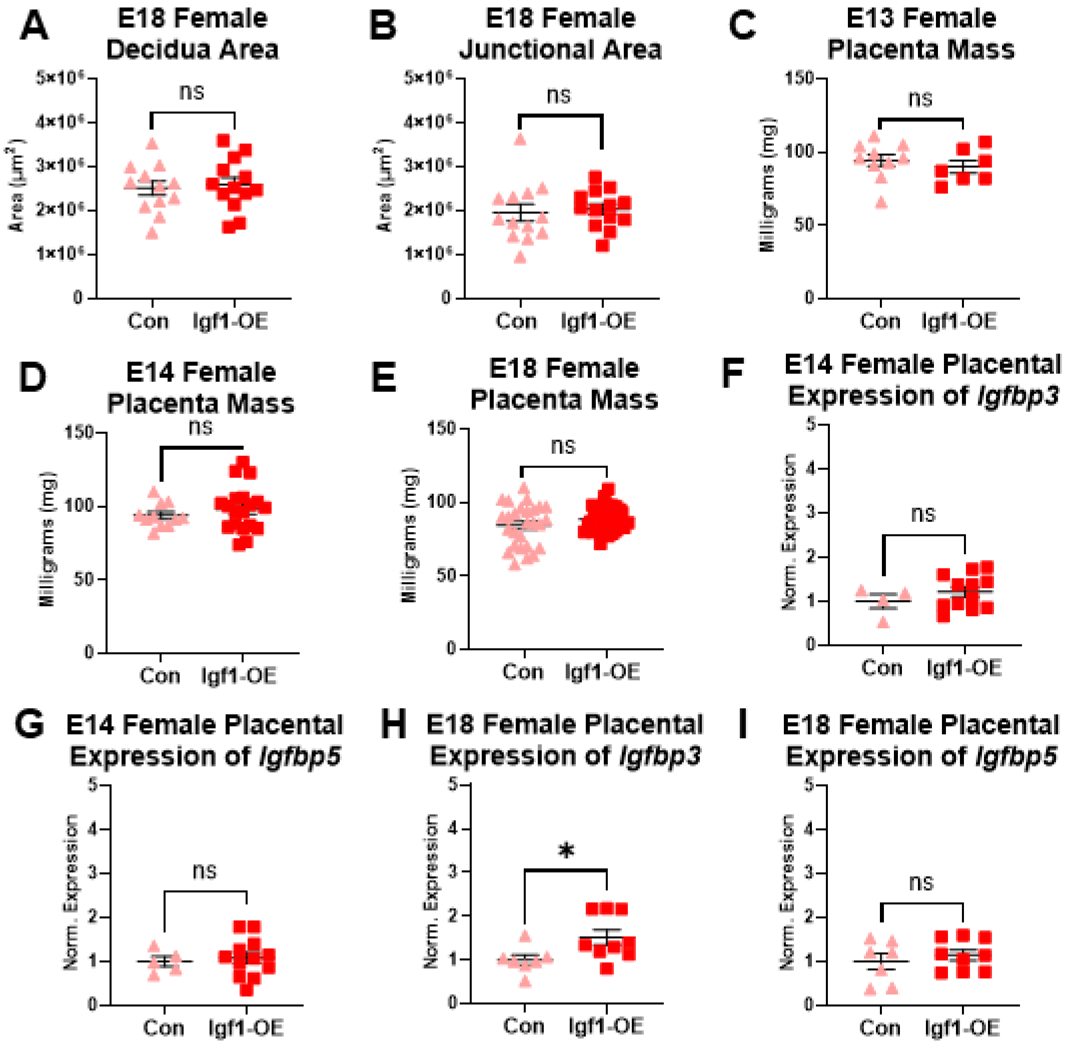


**Supplemental Figure 2: Female placenta structure and gene expression.** Decidua area (A) and junctional zone area (B) in E18 females (n=12-13 per group). Placental mass was observed at E13 (C), E14 (D), and E18 (E) (n=7-35 per group). Placental expression of *Igfbp3* (F) and *Igfbp5* (G) relative to *18s* expression in E14 female placentas (n=4-12 per group). Placental expression of *Igfbp3* (H) and *Igfbp5* expression (I) in E18 females (n=7-9 per group). All graphs show mean and SEM. ns=nonsignificant, *p < 0.05 by linear mixed effects model with litter as a covariate.

#### SUPPLEMENTAL FIGURE 3

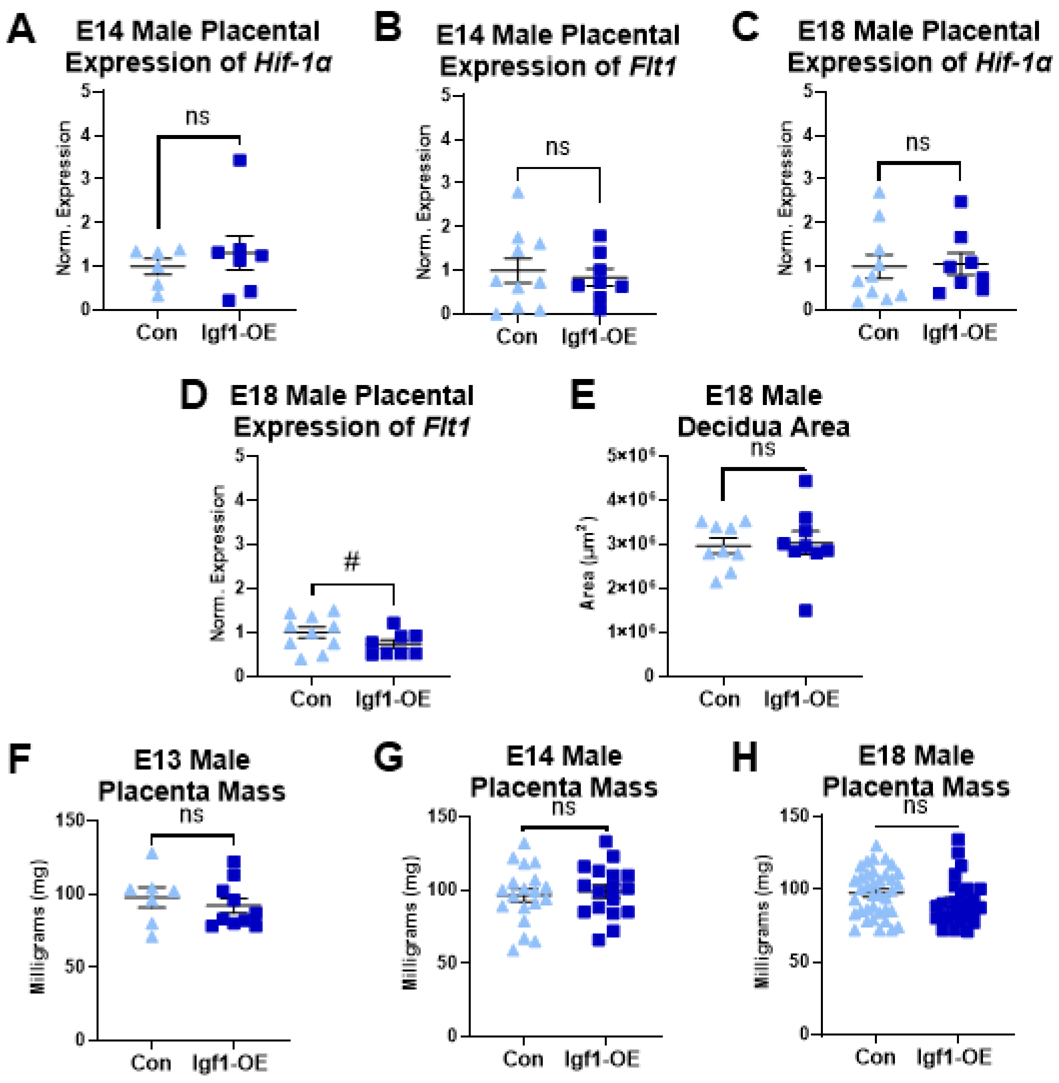


**Supplemental Figure 3: Male placenta structure and gene expression.** E14 male placental expression of *Hif-1α* (A) and *Flt1* (B) (n=6-10 per group). E18 male placental expression of *Hif-1α* (C) and *Flt1* (D) (n=8-10 per group). Decidua area in E18 males (E) (n=9 per group). Male placental mass at E13 (F), E14 (G), and E18 (H) (n=7-37 per group). All graphs show mean and SEM. ns=nonsignificant, #p < 0.1 by linear mixed effects model with litter as a covariate.

**SUPPLEMENTAL FIGURE 4**

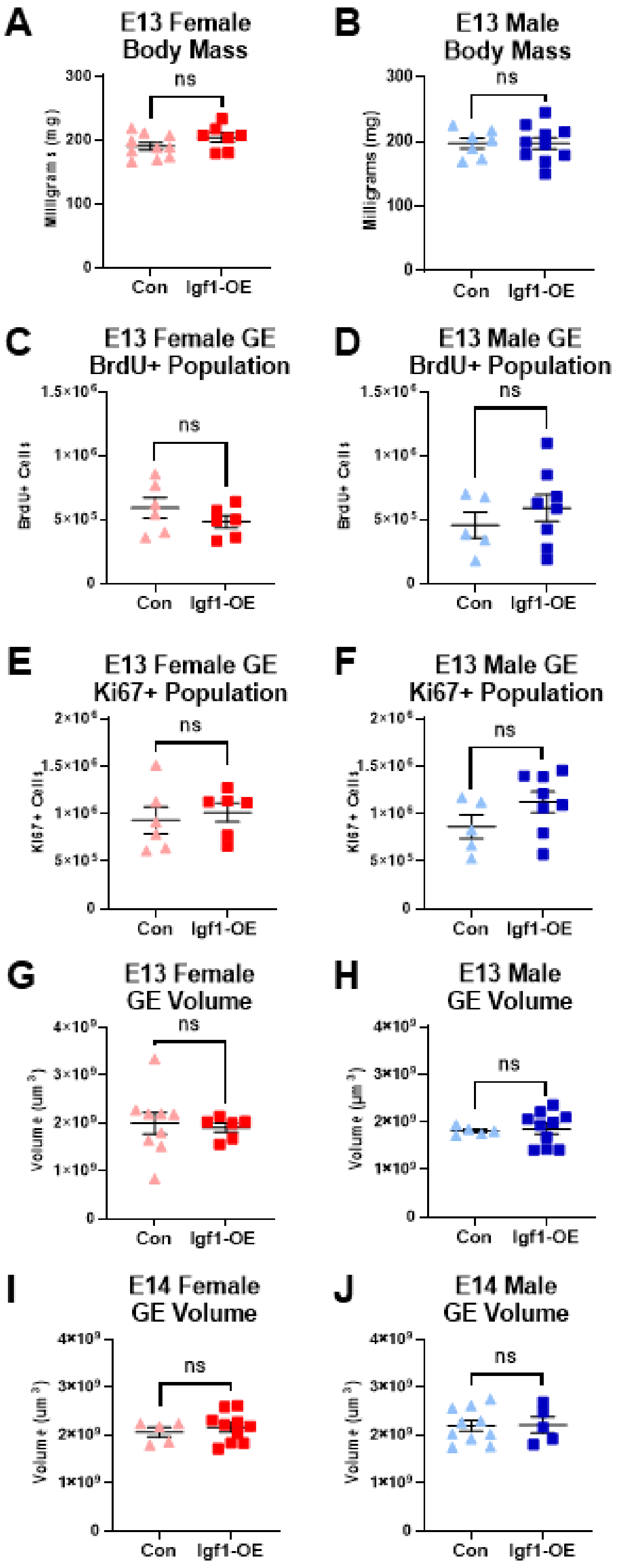

**Supplemental Figure 4: E13 and E14 ganglionic eminence measures.** E13 body mass for females (A) and males (B) (n=7-10 per group). Ganglionic eminence (GE) BrdU+ population at E13 in females (C) and males (D) (n=5-8 per group). GE Ki67+ population at E13 in females (E) and males (F) (n=5-8 per group). GE volume at E13 in females (G) and males (H) (n=5-10 per group). GE volume for E14 females (I) and males (J) (n=5-10 per group). All graphs show mean and SEM. ns=nonsignificant. Analyzed by linear mixed effects model with litter as a covariate.

#### SUPPLEMENTAL FIGURE 5

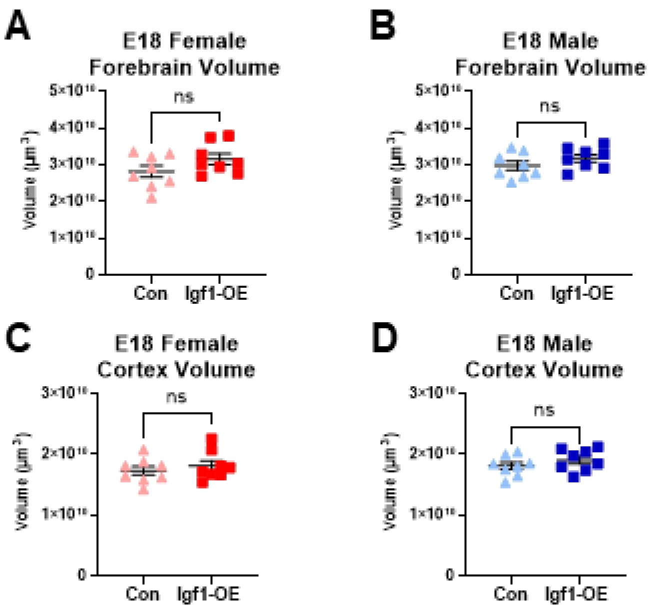


**Supplemental Figure 5: E18 Brain morphology.** E18 forebrain volume in females (A) and males (B) (n=8 per group). E18 cortex volume in females (C) and males (D) (n=8 per group). All graphs show mean and SEM. ns=nonsignificant. Analyzed by linear mixed effects model with litter as a covariate.

#### SUPPLEMENTAL FIGURE 6

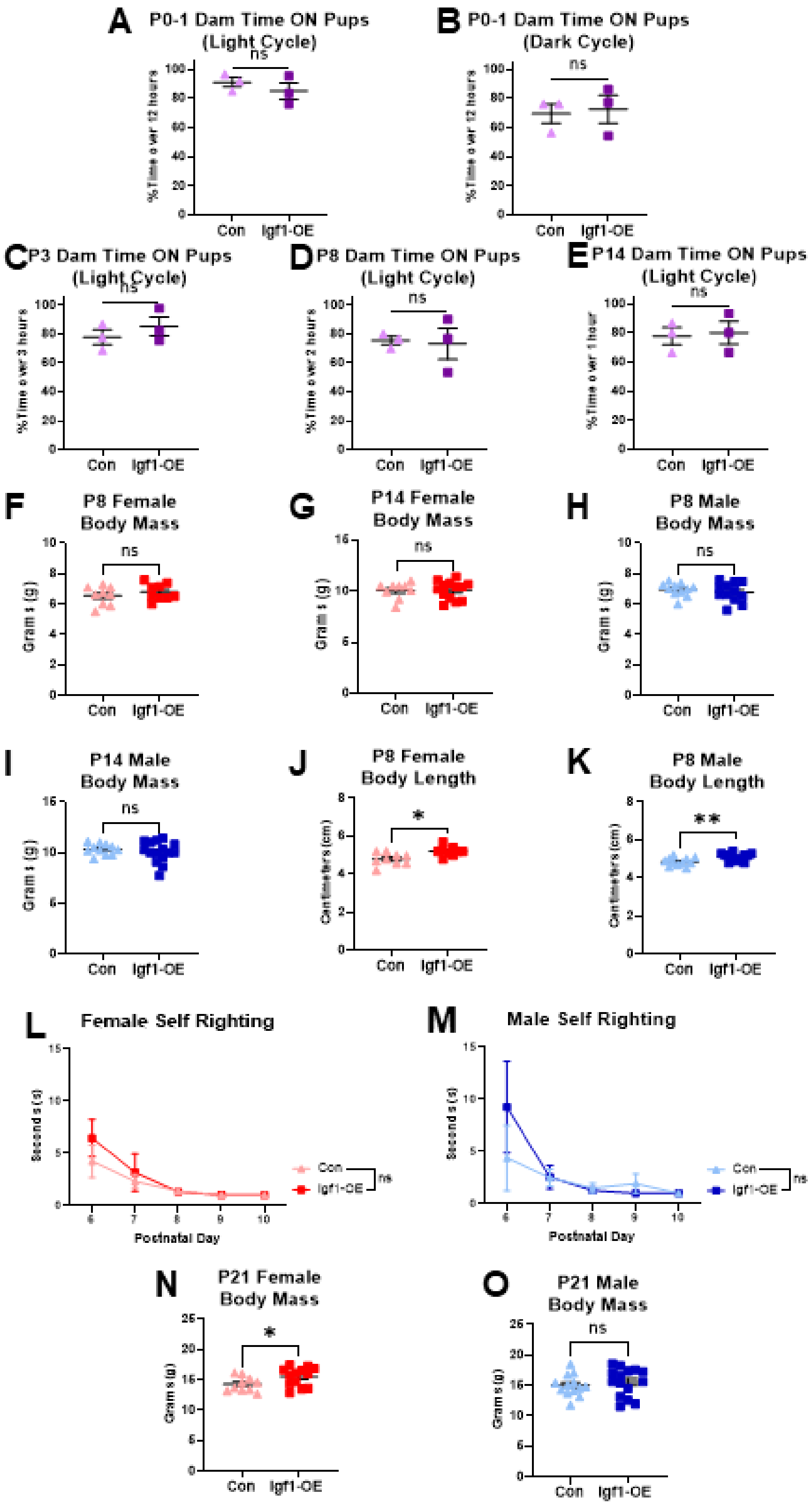

**Supplemental Figure 6: Maternal care and juvenile growth measures.** Maternal care measures across multiple timepoints for dams whose litter received placental-targeted CRISPR manipulation with either all control activation plasmid (Con) or all Igf1-OE plasmid (A-E) (n=3 per group). Body mass for females and males on P8 and P14 (F-I) (n=9-15 per group). Female and male body length at P8 (J,K). (n=9-12 per group). Self-righting testing performed from P6 to P10 results for females or males (L,M) (n=4-12 per timepoint per group). Body mass at weaning on P21 for females and males (N,O) (n=10-15 per group). All graphs show mean and SEM. ns=nonsignificant, *$p < 0.05$ and **$p < 0.01$ by linear mixed effects model with litter as a covariate.

### SUPPLEMENTAL FIGURE 7

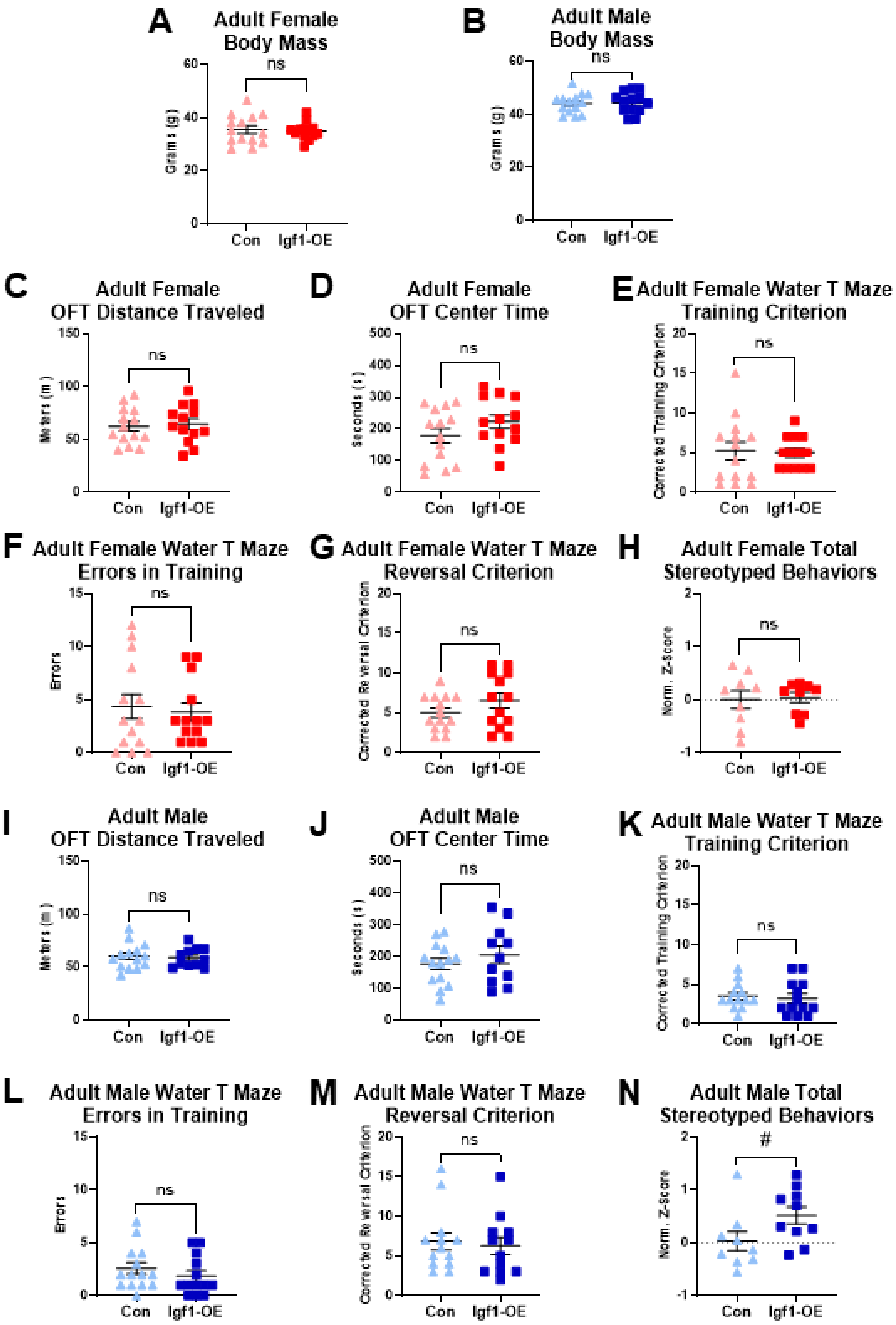

**Supplemental Figure 7: Adult body mass and behavior results.** Adult female and male body mass at time of perfusion (A,B) (n=13-14 per group). Adult female distance traveled in OFT (C), center time in OFT (D), water T maze trials to reach training criterion (E), water T maze errors in training (F), water T maze trials to reach reversal criterion (G), and total stereotyped behaviors (H) (n=9-14 per group). Adult male difference in distance traveled in OFT (I), center time in OFT (J), water T maze trials to reach training criterion (K), water T maze errors in training (L), water T maze trials to reach reversal criterion (M), and total stereotyped behaviors (n=9-14 per group). All graphs show mean and SEM. ns=nonsignificant, #p<0.1 by Welch's t-test.

SUPPLEMENTAL FIGURE 8

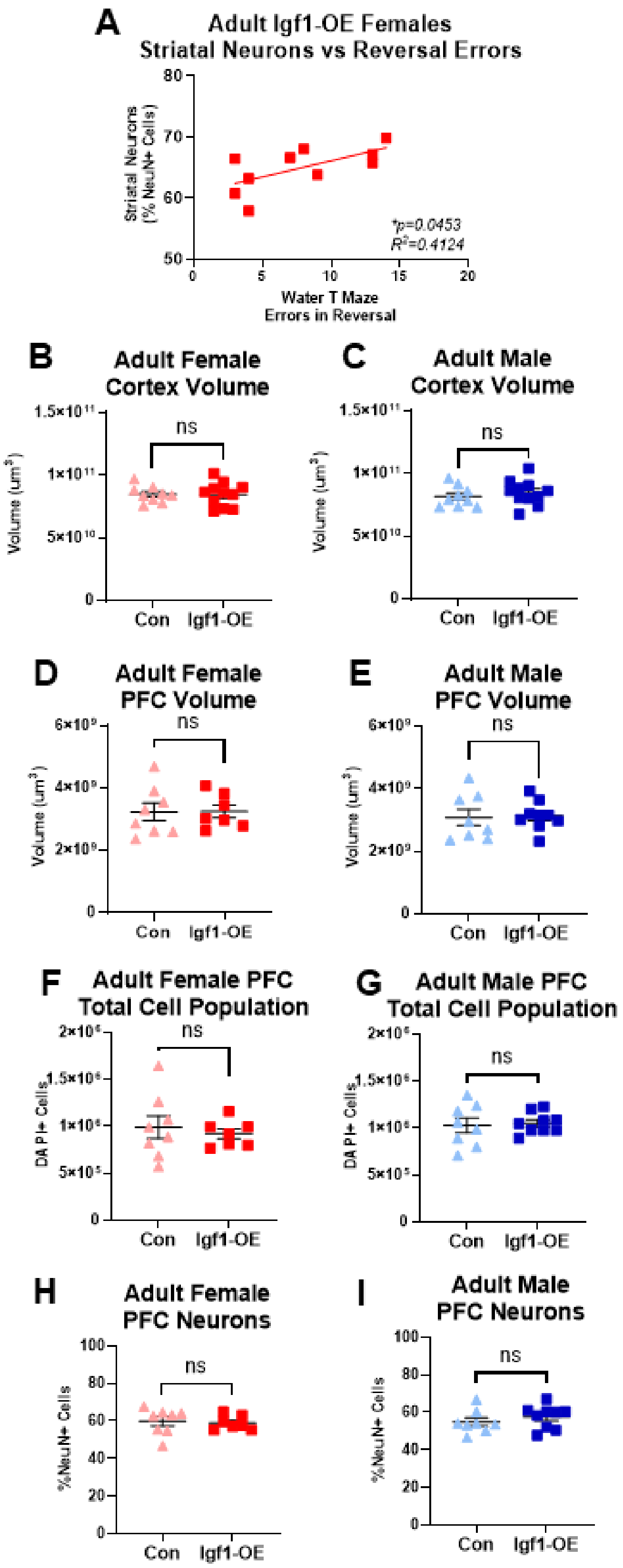

**Supplemental Figure 8: Adult cortex and prefrontal cortex measures.** (A) Correlation between striatal neuron proportion and water T maze errors in reversal for adult Igf1-OE females (n=10 mice). Adult cortex volume for females (B) and males (C) (n=9-11 per group). Adult prefrontal cortex (PFC) volume in females (D) and males (E) (n=9-11 per group). PFC total cell population for females (F) and males (G) (n=7-9 per group). Proportion of neurons in the adult PFC in females (H) and males (I) (n=7-9 per group). All graphs show mean and SEM. ns=nonsignificant, $^{*}p < 0.05$ by Welch's t-test or simple linear regression.
